# Mechanism of neurodegeneration mediated by clonal inflammatory microglia

**DOI:** 10.1101/2024.07.30.605867

**Authors:** Rocio Vicario, Stamatina Fragkogianni, Maria Pokrovskii, Carina Mayer, Estibaliz Lopez-Rodrigo, Yang Hu, Masato Ogishi, Araitz Alberdi, Ann Baako, Oyku Ay, Isabelle Plu, Véronique Sazdovitch, Sebastien Heritier, Fleur Cohen-Aubart, Natalia Shor, Makoto Miyara, Florence Nguyen-Khac, Agnes Viale, Ahmed Idbaih, Zahir Amoura, Marc K. Rosenblum, Haochen Zhang, Elias-Ramzey Karnoub, Palash Sashittal, Akhil Jakatdar, Christine A. Iacobuzio-Donahue, Omar Abdel-Wahab, Viviane Tabar, Nicholas D. Socci, Olivier Elemento, Eli L Diamond, Bertrand Boisson, Jean-Laurent Casanova, Danielle Seilhean, Julien Haroche, Jean Donadieu, Frederic Geissmann

**Author notes:** correspondence to Frederic Geissmann < >, and Julien Haroche. Stamatina Fragkogianni and Maria Pokrovskii equally contributed.

## Abstract

Langerhans cell Histiocytosis (LCH) and Erdheim-Chester disease (ECD) are clonal myeloid disorders, associated with MAP-Kinase activating mutations and an increased risk of neurodegeneration. Surprisingly, we found pervasive PU.1^+^ microglia mutant clones across the brain of LCH and ECD patients with and without neurological symptoms, associated with microgliosis, reactive astrocytosis, and neuronal loss. The disease predominated in the grey nuclei of the rhombencephalon, a topography attributable to a local proliferative advantage of mutant microglia. Presence of clinical symptoms was associated with a longer evolution of the disease and a larger size of PU.1^+^ clones (p= 0.0003). Genetic lineage tracing of PU.1^+^ clones suggest a resident macrophage lineage or a bone marrow precursor origin depending on patients. Finally, a CSF1R-inhibitor depleted mutant microglia and limited neuronal loss in mice suggesting an alternative to MAPK inhibitors. These studies characterize a progressive neurodegenerative disease, caused by clonal proliferation of inflammatory microglia (CPIM), with a decade(s)-long preclinical stage of incipient disease that represent a therapeutic window for prevention of neuronal death.

## Introduction

Langerhans cell histiocytosis (LCH)^1^ and Erdheim-Chester disease (ECD) ^2,3^, are rare clonal myeloid disorders, with an estimated incidence of ∼5 cases per million people per year ^4^. LCH and ECD, sometimes co-diagnosed in the same patients ^5^, are both associated with a high prevalence of the BRAF^V600E^ mutation^1–3,6–8^ as well as an increased risk of neurodegenerative disease (neuro-histiocytosis) ^1,3,9–11^. LCH and ECD were grouped under the new name of “L-histiocytosis” ^5^, but LCH is predominantly a pediatric disease, while ECD is diagnosed in adults and their clinical presentation typically differs^5^. However, in both LCH and ECD patients, neuro-histiocytosis is characterized by progressive symmetric cerebellar syndrome, tetra-pyramidal syndrome with or without motor deficits, pseudobulbar palsy, and cognitive and behavioral impairment ^10–14^. The natural history of neurodegeneration is not precisely documented, but it can occur decades after the original diagnosis of histiocytosis, in patients considered « in remission » ^1,10,12^, albeit it can also be the initial presentation of the disease ^3^. Brain MRI abnormalities, frequently non-specific, include T1 hypersignal of deep pons and cerebellar grey nuclei, demyelination and atrophy located preferentially to posterior fossa ^3,12–16^. Full brain pathological examination was only reported in 3 cases in the medical literature which reported neuronal loss, gliosis, and demyelination^17^. The mechanism of neurodegeneration is poorly understood^12,13,17^. The cell of origin is also an outstanding question. Mutant bone marrow hematopoietic progenitors were proposed to invade the brain in some cases ^18–21^. In addition, mosaic expression of BRAF^V600E^ in the resident macrophage lineage ^22–24^ in a mouse model was associated with proliferation of BRAF^V600E^ macrophages in the lung, liver and microglia in the brain, and with a neurodegenerative disease in the absence of bone marrow-derived clones^25^. Given the need to understand the underlying mechanisms of neurodegeneration in LCH and ECD for the identification of novel therapeutic strategies and targets, we undertook a comprehensive and systematic analysis of the brains from a series of LCH and ECD patients.

## Results

### Pervasive BRAF^V600E^ PU.1^+^ clones in the brain of LCH and ECD patients

We studied 8 consecutive patients (**Table 1**) diagnosed with pediatric onset LCH (n=2), adult onset mixed LCH/ED (n=2), or adult onset ECD (n=4) based on BRAF^V600E^ positive lesions, for whom post-mortem whole brain (in 7 cases) or brain biopsy (n=1, Patient#2) and blood or bone marrow samples were obtained after informed consent for the purpose of this study (**Table S1**). Four patients were diagnosed with neuro-histiocytosis in the course of the disease and the other 4 were free of neurological symptoms (**Table 1 and Extended Data Patients**). Brain and blood samples from 35 neuro-typical individuals without histiocytosis (**Table S1**) were also studied as controls. Blood or Bone Marrow (BM) samples and nuclei suspensions from frozen brain tissue sampled from the frontal cortex to the spinal cord, and FACS sorted into PU.1^+^ myeloid nuclei, NeuN^+^ neuronal nuclei^26^, and PU.1^-^ NeuN^-^ (DN) stromal nuclei (**Figure 1A and Figure S1A**), were subjected to targeted deep sequencing (^27^, **see Methods**, **Table S2, S3**) at an average depth of ∼1100X (**Figure S1B**), and to droplet digital PCR (ddPCR) (**Table S3**). BRAFc.1799T>A (corresponding to BRAF^V600E^) was the variant most frequently detected, present in multiple PU.1^+^ brain samples, from all histiocytosis patients (**Figure 1B and Table S3**). BRAF^V600E^ detection was confirmed by ddPCR (**Table S3**). In contrast, BRAF^V600E^ was not detected in NeuN^+^ and DN samples from histiocytosis patients and in NeuN^+^, PU.1^+^ and DN samples from controls (**Figure 1C, Figure S1C and Table S3**). These results indicated that the BRAF^V600E^ mutation, although specific for patients in comparison to controls, was -surprisingly-pervasive among PU.1+ nuclei across the brain, even in the absence of clinical neurological symptoms.

**Figure 1.**
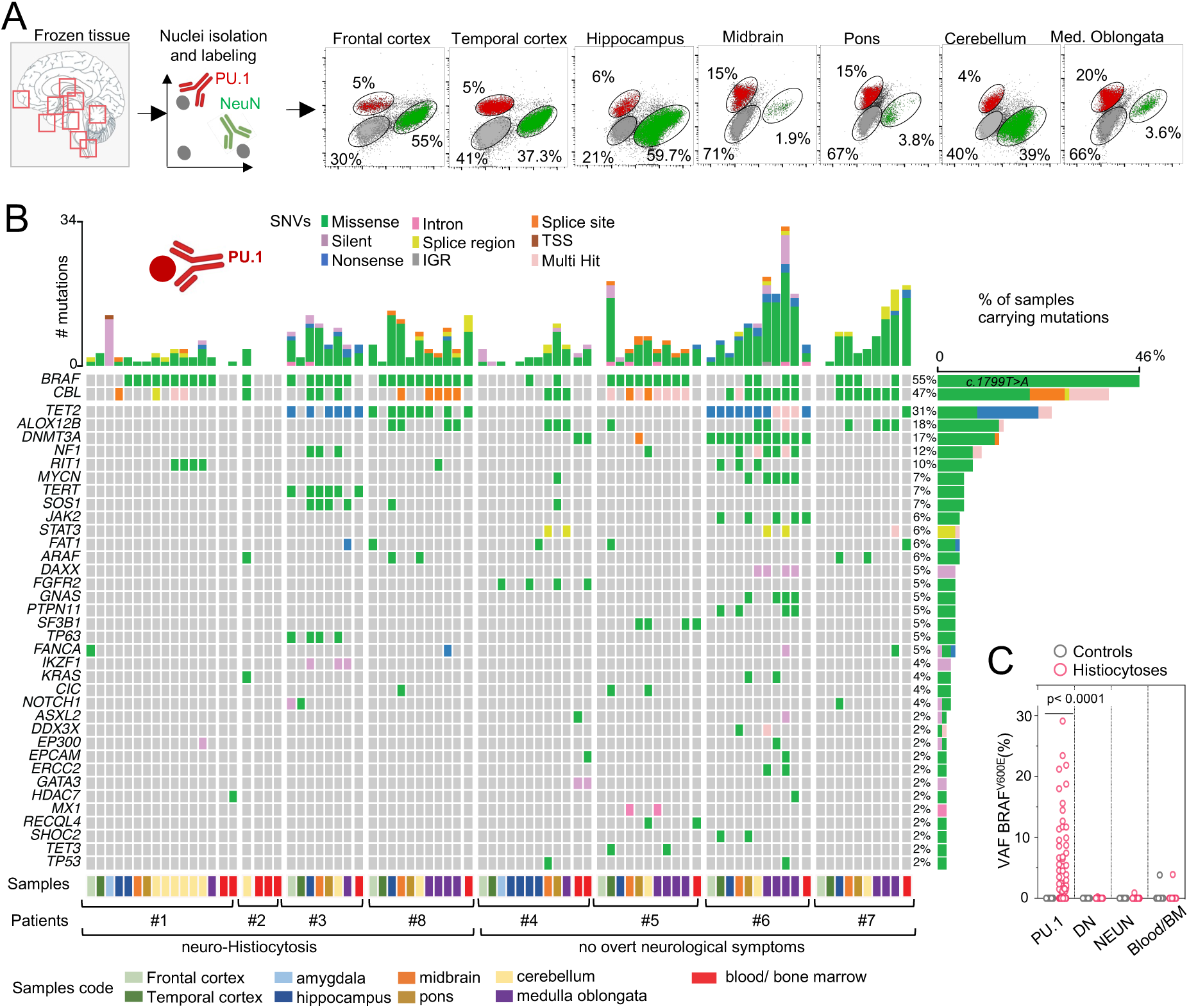
Detection of mutations in brain and matching blood or bone marrow from histiocytosis patients. **(A)** Left, schematic of post-mortem brain samples obtained from patients. Right, representative flow cytometry dot-plots of brain nuclei from patient #1 and labeled with anti-NeuN and anti-PU.1 antibodies (% of total). **(B)** Oncoplot represents mutated genes (with 4 or more mutant reads), number of mutations per sample and % of samples carrying mutations in PU.1+ samples (n=71) and matching blood or bone marrow samples (BM) (n=12) from Histiocytosis patients (n=8). **(C)** Variant allelic frequency (VAF, %, HemePACT) for BRAFc.1799T>A (V600E) in PU.1+, NeuN+, DN (PU.1-,NeuN-) and blood or bone marrow samples from Histiocytosis patients (71 brain samples from 8 patients), and controls (104 samples from n=35). Each dot represents a sample. Statistics: p-values are calculated with unpaired two-tailed Mann-Whitney U test.

**Table 1.** Characteristics of L-Histiocytosis patients. . DX: Diagnosis (L: Langerhans Cell Histiocytosis; E: Erdheim-Chester Disease); BM: Bone marrow; na: not available; Neuro.: clinical diagnosis of Neuro-histiocytosis. Delay: years from initial diagnosis and Neuro-histiocytosis diagnosis. y: years.

|  | Dg | Age | Sex | BRAF<sup>V600E</sup> | Tissues analyzed | Age at DX LCH/ECD | Extra-neurological symptoms of histiocytosis | Age at DX of Neuro-Histio | Delay between diagnosis of histio and Neuro-histio | Brain MRI | Other diseases | Cause of death |
| --- | --- | --- | --- | --- | --- | --- | --- | --- | --- | --- | --- | --- |
| #1 | LCH | 21 | M | + | Brain, blood | 0.75 | Skin, bone | 13 | 12 y | + | none | Neuro-Histio |
| #2 | LCH | 26 | F | + | Cerebellar biopsy, blood | 0.5 | Skin, bone | 16 | 15 y | + | none | alive with severe Neuro-Histio |
| #3 | LCH/ECD | 78 | M | + | Brain, bone marrow | 52 | Skin, bone, heart, lung | 77 | 25 y | + | Clonal hematopoiesis (TET2) | Neuro-Histio |
| #8 | ECD | 69 | F | + | Brain, bone marrow | 58 | Bone, Retro-Peritoneal, heart | 66 | 8 y | + | Metastatic breast Cancer<br>Clonal hematopoiesis (TET2, BRAF) | Breast Cancer |
| #4 | ECD | 66 | M | + | Brain, bone marrow | 59 | Skin, bone, heart, lung, Retro-Peritoneal | - |  | - | Pancreatic cancer<br>Clonal hematopoiesis (DNMT3) | Pancreatic Cancer |
| #5 | LCH/ECD | 78 | F | + | Brain, bone marrow | 75 | Skin, bone, heart, lung, liver | - |  | - | none | Hepatic insufficiency |
| #6 | ECD | 61 | M | + | Brain, bone marrow | 59 | Skin, bone | - |  | nd | Clonal hematopoiesis (TET2/JAK2/DNMT3)<br>T cell lymphoma | T cell lymphoma |
| #7 | ECD | 89 | M | + | Brain, bone marrow | 85 | Bone, Retro-Peritoneal | - |  | - | Kidney cancer<br>Prostate cancer<br>Clonal hematopoiesis (TET2)<br>Stroke | Stroke, sepsis |

### Overt and incipient neurodegeneration in histiocytosis patients

Neuropathological analysis of corresponding brain samples (**see Extended Data Patients**) indeed revealed the presence of histological lesions in all patients, including the 4 patients without clinical neurodegeneration (**Figure 2A**, *red and purple arrows*). Histological lesions were found in the anatomical areas where the BRAF^V600E^ mutation was identified (**Figure 2A**, *red arrows*) and consisted in focal, non-systematized, areas of microglial (IBA1+) and astrocyte (GFAP) activation, and neuronal loss in the grey matter (**Figure 2B)** and axonal spheroids in the white matter. The pons, cerebellum, and hippocampus were the most frequently affected, while the cortex was rarely involved (**Figure 2A, B and Figure S2A**). Differential gene expression and pathway analyses of whole-tissue RNAseq from patients and control brains showed upregulation of inflammatory and phagocytic signatures including complement, IL1, and phagocytic receptors, mainly driven by the patient’s brainstem and cerebellum samples, and down-regulation of genes associated with neuronal and synaptic activity (**Figure 2C, D and Table S4**). Of note, a posteriori review of available brain MRIs showed the presence of nonspecific hyperintense signals in the dentate nuclei of 2 of 3 patients without clinical symptoms, reminiscent of nonspecific hyperintense signals found in the pons, dentate nuclei, and cerebellum of patients with neurological symptoms (**Figure S2B**).

**Figure 2.**
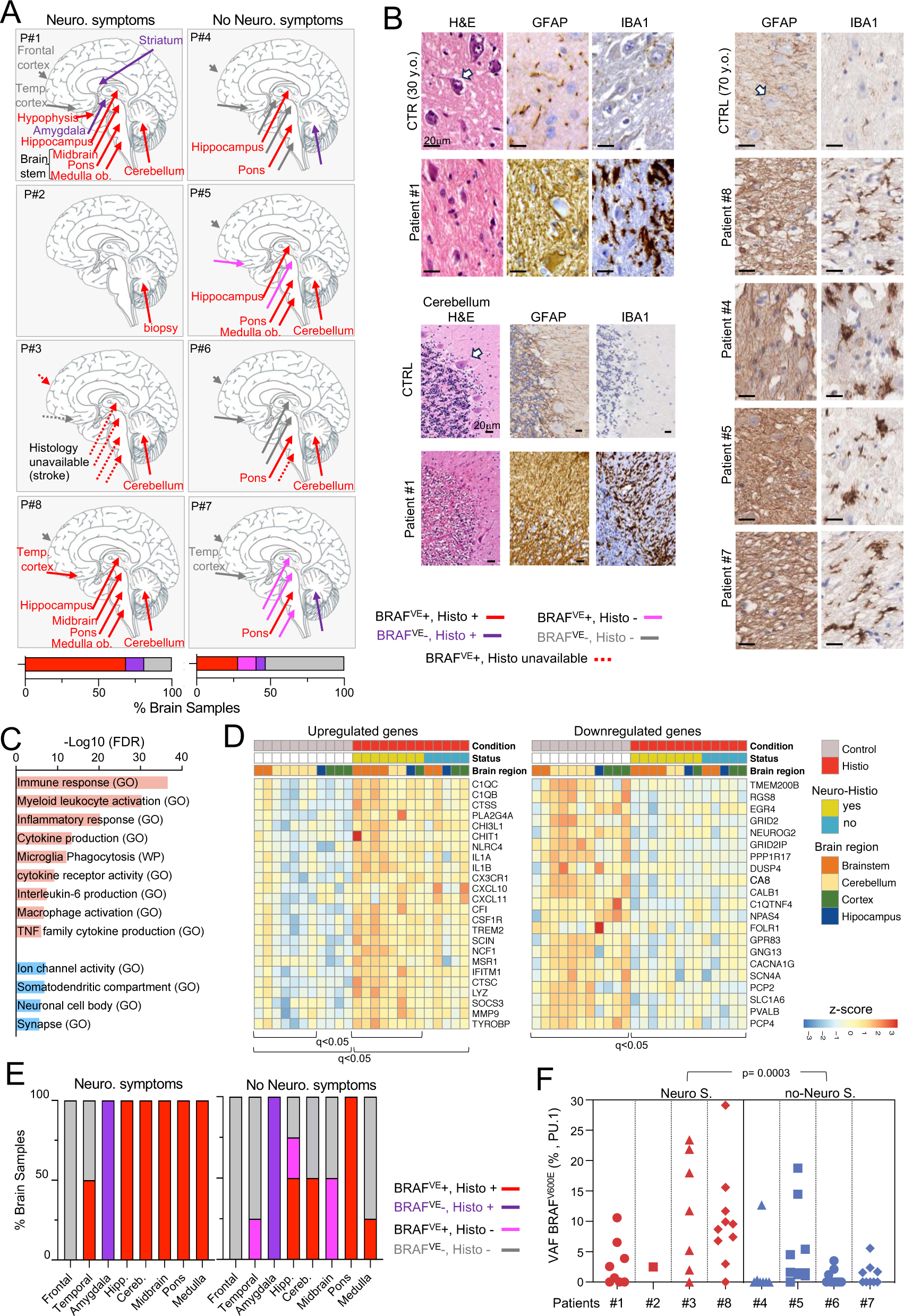
Histological and molecular analysis of the brain of histiocytosis patients. **(A)** Schematic of the brains of the 8 patients annotated for the detection of BRAFc.1799T>A (V600E) and/or of histological signs of neurodegeneration (Histo+). Bar graphs represents the proportion of tested brain samples positive for BRAFc.1799T>A (V600E) by HemePACT and/or histological signs of neurodegeneration among patients with (left) or without (right) neurological symptoms**. (B)**. Left, representative H&E, IBA1 (microglia marker) and GFAP (astrocyte marker) of pons and cerebellum from patient #1 and an age-match control for comparison. Right, representative IBA1 and GFAP of pons from patient #8, #4, #5, #7 and an age-match control for comparison. Arrows indicate neuron nuclei. **(C)** Pathway enrichment among differentially expressed genes (DEG, red: upregulated genes, blue: downregulated genes) by RNAseq analysis (FDR <0.05, log2FC >= or <= 1.5/-1.5) of whole brain of Histiocytosis patients (n=13) and controls (n=11) using g:profiler webtool. Pathways are selected based on FDR <= 0.05 and ordered by significance. **(D)** Hierarchical clustering of DEG (log2FC >= 1.5, log2FC <= −1.5, FDR <0.05) between brain samples from Histiocytoses (n=13) and controls (n=11). Expression values are Z score transformed. **(E)** Bar graphs represents the proportion of tested brain samples positive for BRAFc.1799T>A (V600E) by HemePACT and/or histological signs of neurodegeneration among patients with (left) or without (right) neurological symptoms. **(F)** Variant allelic frequency (VAF, %, HemePACT)) for BRAFc.1799T>A (V600E) in PU.1+, samples from patients with (red) and without (blue) neurological symptoms. Each symbol represents a patient. Statistics: p-value is calculated using a mixed-effects linear regression model (see methods).

These data indicated that the patients with pediatric onset LCH and adult onset ECD presented with qualitatively similar molecular and cellular signs of neurodegeneration, whether they presented or not with neurological symptoms, and therefore that patients without clinical symptoms presented with ‘incipient’ BRAF^V600E^-associated neurodegeneration.

### Preclinical neurodegeneration characterized by BRAF^V600E^-mediated clonal expansion

As shown above, areas of histological neuroinflammation and neuronal loss overlapped with molecular detection of the BRAF^V600E^ variant in the same regions (red arrows, **Figure 2A**), however, this overlap was more consistent in patients with clinical neuro-histiocytosis than in patients without neurological symptoms (**Figure 2A, E and Figure S2C**). In addition, analysis by 2 neuropathologists determined that histological damage in the pons was more intense in patients with clinical symptoms (**Figure S2D**). More quantitatively, a mixed-effects linear regression model analysis (see Methods) showed that the average size of the BRAF^V600E^ clone was a significant predictor of the presence of neurological symptoms, despite the small number of patients (p=3e^-4^, **Figure 2F**). Of note, the delay between the initial diagnosis of LCH or ECD and the diagnosis of neuro-histiocytosis was of 8 to 25 years, while the 4 patients without clinical neurological symptoms died from other causes 2 to 7 years after diagnosis of ECD (**Table 1**).

These results altogether demonstrate that patients develop molecular and histological features of incipient neurodegeneration within a few years from the initial diagnosis of histiocytosis and possibly decade(s) before development of clinical symptoms, while clinical symptoms and the severity of histological changes correlate with the size of the BRAF clone, suggesting progressive damage as mutant clones expand.

### Preferential proliferation of BRAF^V600E^ microglial clones in the mammalian rhombencephalon

As shown above, detection of BRAF^V600E^ clones, neuronal death, astrogliosis, and microgliosis, and the neuroinflammatory signature predominated in the patients’ hippocampus, brainstem and cerebellum (rhombencephalon) (**Figure 2A**), in accordance with the classical cerebellar syndrome, pseudobulbar palsy, and cognitive and behavioral impairment reported in neuro-histiocytosis ^3,11–13^. To explore the mechanisms involved in the anatomical topography of the clonal process, we performed an analysis of the allelic frequency (AF) distribution of the BRAF^V600E^ variant as a function of the location of samples, from the frontal cortex to the medulla oblongata. Results showed that the size of BRAF^V600E^ clones by HemePACT as well as by ddPCR increases along a rostro-caudal gradient (r =0.75, p=0.005, and r= 0.7, p= 0.0089 respectively, **Figure 3A and Figure S3A**). Average AF of BRAF^V600E^ clones along a rostro-caudal gradient was increased in patients with as well as without neurological symptoms (r= 0.67, p: 0.01, r= 0.8, p= 0.002, **Figure 3A**).

**Figure 3.**
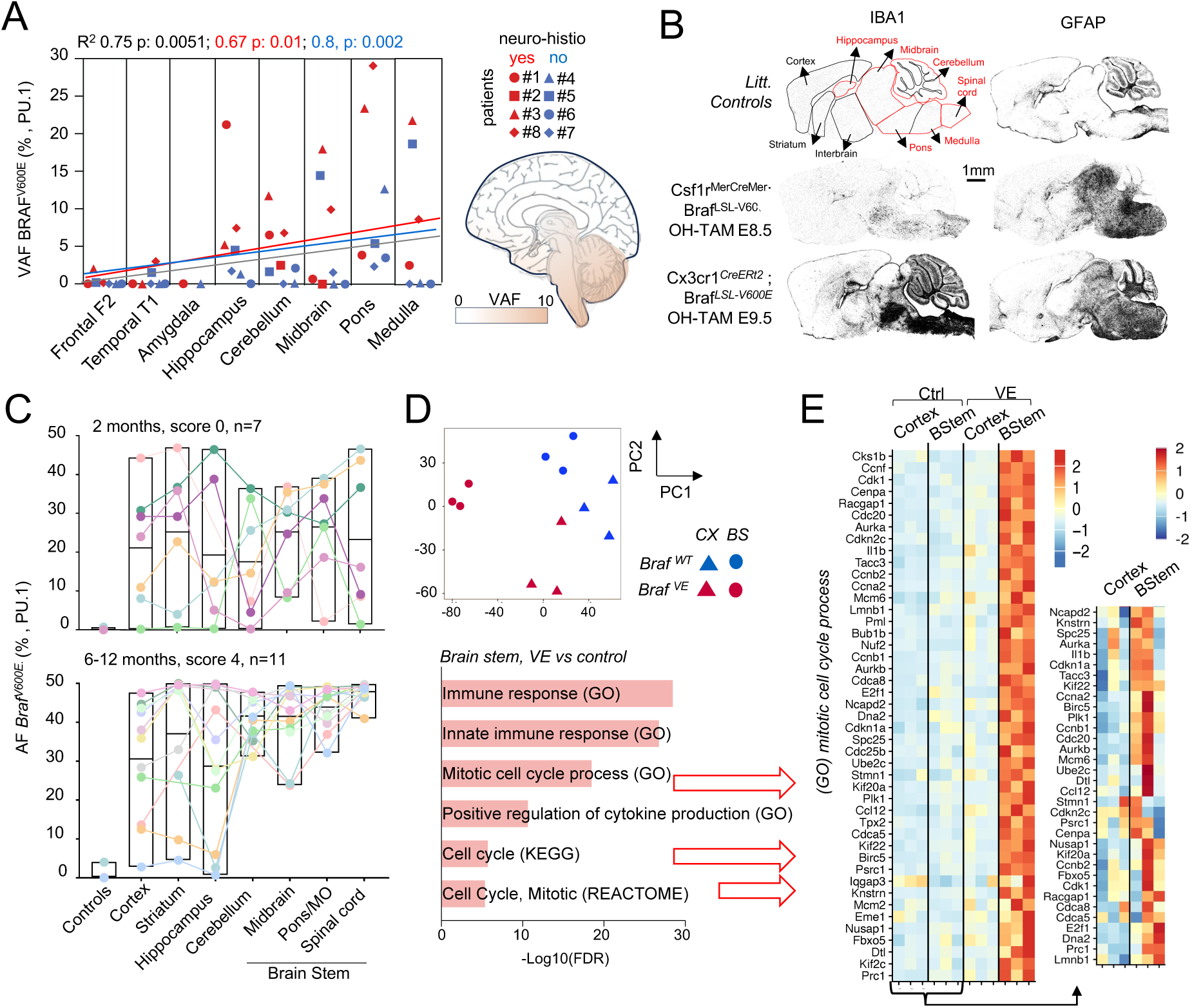
Analysis of mutant microglia across brain regions in human and mouse models of neuro-histiocytosis. **(A)** Variant allelic frequency (VAF, %, by HemePACT) of BRAFc.1799T>A (V600E) in PU.1+ nuclei from histiocytosis patients across brain regions (n=8, patients with neuro-histiocytosis are color-coded in red, patients without a diagnosis of neuro-histiocytosis are color-coded in blue). The fitted line, R-squared and corresponding p value were calculated by simple linear regression by assigning numbers from 1-8 to each brain region from along a rostro caudal axis. Gray line: all patients. Red line: patients with neuro-histiocytosis. Blue line: patients without neuro-histiocytosis. **(B)** Representative mouse sagittal midline brain sections from 6 months old *Csf1r^MerCreMer^; Braf^LSL-V600E^* mice (pulsed with OH-TAM at E8.5*), Cx3cr1^CreERt2^ ; Braf^LSL-V600E^* (pulsed with OH-TAM at E9.5) and littermate controls stained with anti-IBA1 or anti-GFAP, Scale bar 1000uM. **(C)** Allelic frequency of the Braf^V600E^ allele in microglia purified from dissected brain regions from 2 months old, and at 6-12 months-old analyzed by droplet digital PCR (ddPCR). Dots and colored lines represent individual mice, boxes represent variance, with line at mean. **(D)** RNAseq analysis performed in FACS-isolated microglia from cortex and brainstem from 2-month-old *Cx3cr1^CreERT2^ Braf^LSL-V600E^* mice (n=3), and littermates (n=3) pulsed with OH-TAM at E8.5. Top, principal component analysis (PCA). Bottom, pathway analysis of significantly upregulated genes (FDR <0.05, log2FC >= 1.5) in microglia from old *Cx3cr1^CreERT2^ Braf^LSL-V600E^* versus littermate control using g:profiler webtool. Pathways are selected based on FDR <= 0.05 and ordered by significance. **(E)** Hierarchical clustering of DEG from ‘mitotic cell cycle process’ (GO:1903047, left) from analysis in D.

This anatomical distribution of microglial clones and brain damage could correspond in theory to preferential engraftment of circulating clones and/or to a local survival or proliferative advantage of mutant clones in the hindbrain. Mouse models of neuro-histiocytosis, generated by mosaic targeting of a Braf^V600E^ allele in embryonic resident macrophages, result in focal proliferation and activation of macrophages across several organs, including microglia, followed by paralysis and neurodegeneration after ∼6 month of life (^25^, **Figure 3B and Figure S3B-D**). We therefore measured Braf^V600E^ allelic frequency in microglia from experimental mice and control littermates, along the brain rostro-caudal axis of over time. Interestingly, Braf^V600E^ microglia was randomly distributed throughout the brain in young mice, at various allelic frequencies but later selectively accumulated in the rhombencephalon (**Figure 3C**) phenocopying the results from patients and suggesting a local survival or proliferative advantage of the mutant clones.

Pathway analysis of differentially expressed genes in RNAseq from FACS-sorted microglia from the cortex and brainstem of mice and control littermates showed a vigorous and brainstem-specific microglia proliferative response associated with an inflammatory signature and the absence of cellular senescence (**Figure 3D, E and Figure S4**). As previously shown^28^ the proliferative activity of wild-type microglia is higher in brainstem than the cortex however, this effect was exacerbated in Braf^V600E^ microglia (**Figure 3E**). It is of note that enforced expression of BrafF^V600E^ at high allelic frequency in both cortex and rhombencephalon resulted instead in microglia and astrocyte activation in both cortex and rhombencephalon (**Figure S5**), suggesting that cortex microglia may not *per se* be refractory to activation by Braf^V600E^. Altogether, analysis of mouse models strongly suggests that the preferential accumulation of BRAF^V600E^ microglial clones in the mammalian rhombencephalon is driven, at least in part, by a local proliferative advantage of microglia amplified by the BRAF^V600E^ mutation.

### Natural history of the patients’ PU.1^+^ BRAF^V600E^ clones

Single nuclei (sn)-RNAseq indicated that PU.1^+^ nuclei include ∼93% microglia-like nuclei (^29^ **see Methods**), but does not allow to distinguish a resident or bone marrow origin of these microglial-like cells^29^. We therefore used a genetic bar-coding approach to investigate the putative origins of the patients PU.1^+^ BRAF^V600E^ clones. Analysis of all single nucleotide variants (SNV) identified by deep sequencing showed that the clonal diversity of brain PU.1+ nuclei is independent of blood, *i.e.* brain PU.1+ SNV were rarely detectable in corresponding blood or bone marrow samples, and vice-versa blood/BM SNVs were rarely detected in the brain (**Figure 1B and Figure 4A,B**). These results are consistent with the local maintenance and clonal diversification of brain resident microglia^22,25,30–32^. Nevertheless, the BRAF^V600E^ variant was detected in the in the bone marrow and blood from two ECD patient (#8 and #3) and clonal hematopoiesis (CH) ^33,34^ carrying TET2 or DNMT3A variants was identified in the blood/BM of 5 patients, also detected in the brain of 3 of them (#8, #3 and #6) (**Figure 1B**, **Figure 4C**). We therefore investigated lineage relationship between BRAF and CH clones in the brain and blood/bone marrow in our series of patients.

**Figure 4.**
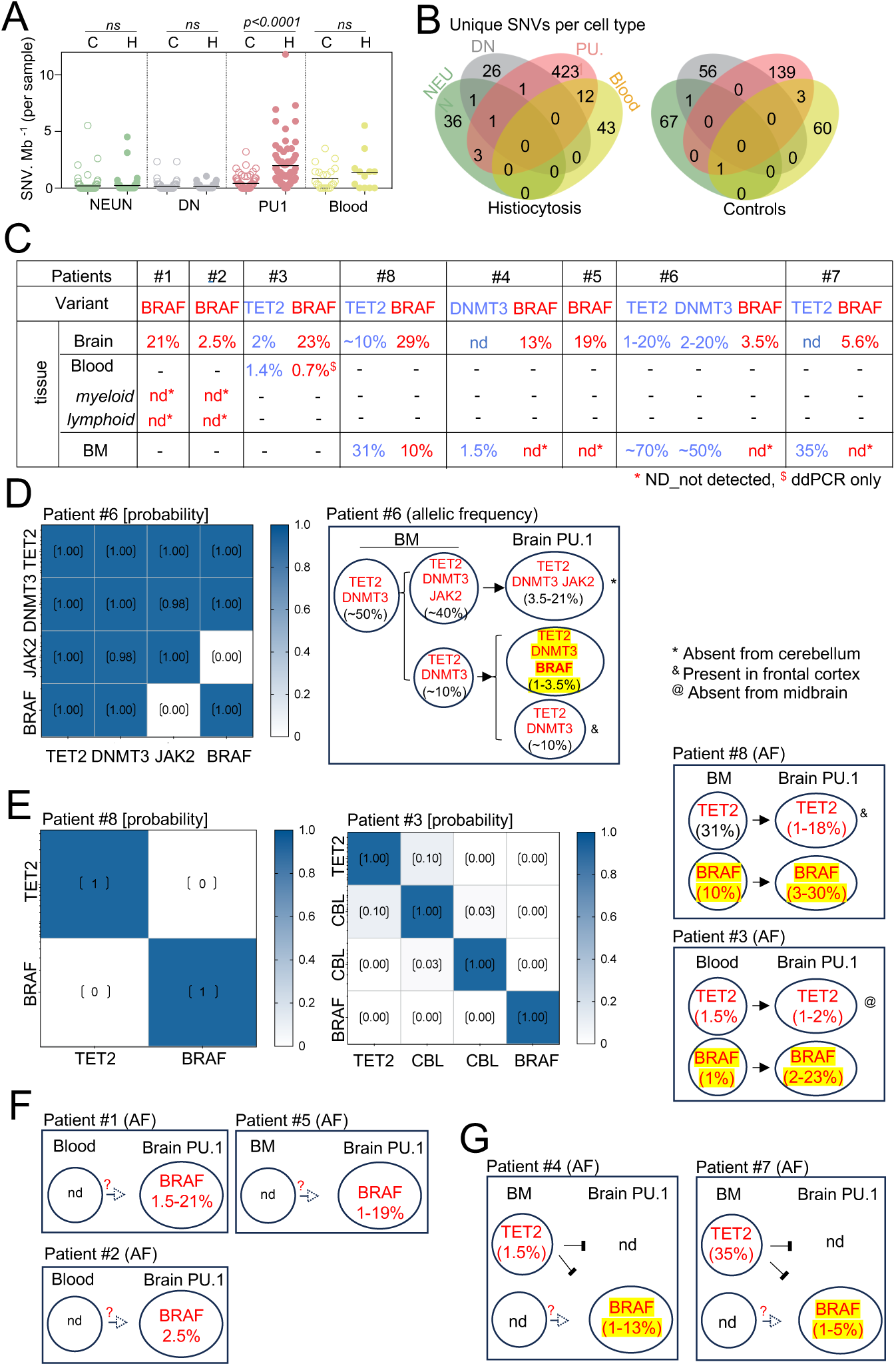
Natural history of BRAF^V600E^ clone. **(A)** Mutational load in NeuN, DN (double negative, NeuN-, PU.1-), PU.1 and Blood/Bone Marrow from control and Histiocytosis patients. Statistics: p-values are calculated with unpaired two-tailed Mann-Whitney U test. **(B)** Venn diagrams represent the repartition per cell type of single-nucleotide variations (SNVs) identified in NeuN^+^, PU.1^+^, DN and matching blood in samples from 8 histiocytosis patients (NeuN, n=62; DN, n=69; PU.1, n=71; Blood/BM, n=12) and 35 control individuals (NeuN, n=107; DN, n=108; PU.1, n=107; Blood/BM, n=22). **(C)** Variant allelic frequency (VAF, %, HemePACT) for BRAF c.1799T>A (V600E) (red), TET2 and DNMT3A variants (blue) in brain PU.1+nuclei and blood/bone marrow nd: not detected (*mean depth 5600x, see Figure S5). For patient #1 and #2 myeloid (HLA-DR+, Lin-) and lymphoid (Lin+) cells were flow-sorted. **(D)** Left, mutual exclusivity analysis of mutations found by single-cell genotyping (Tapestri) of PU.1+ cells isolated from Pons and Cerebellum from patient #6. The number represents the probability that two mutations are mutually exclusive in single cells by random chance. The smaller the probability, the more likely they are in different cell populations. Right, plot depicts probable origin of the BRAFV600E and CH clones in patients #6. Numbers show the range of allelic frequency in different brain samples for each mutation. **(E)** Left, mutual exclusivity analysis of mutations found by single-cell genotyping (Tapestri) of PU.1+ cells isolated from Pons from patient #8 and Cerebellum from patient #3. The number represents the probability that two mutations are mutually exclusive in single cells by random chance. The smaller the probability, the more likely they are in different cell populations. Right, plot depicts probable origin of the BRAFV600E and CH clones in patients #8 and #3. Numbers show the range of allelic frequency in different brain samples for each mutation. **(F)** Plot depicts probable origin of the BRAFV600E clones in patients #1, #2 and #5 based on sequencing data. Numbers show the range of allelic frequency in different brain samples for each mutation. **(G)** Plot depicts probable origin of the BRAFV600E clones and CH clones in patients #4 and #7 based on sequencing data. Numbers show the range of allelic frequency in different brain samples for each mutation.

Single nuclei genotyping (Tapestri) of brain PU.1+ nuclei from patient #6, which presented with clonal hematopoiesis (TET2 and DNMT3 variants at AF∼50%) with a JAK2 variant subclone at 40% AF, as well as a brain specific BRAF^V600E^ clones (**Figure 1B**, **Figure 4C, Table S3**), showed that the BRAF^V600E^ nuclei present in the patient’s brain (at AF 3.5%) also carried the TET2 and DNMT3 variants (**Figure 4D**). It is of note that the PU.1^+^ TET2/ DNMT3/JAK2 variant was absent from the cerebellum (**Table S3**). Therefore, it is possible to conclude that a bone-marrow derived brain PU.1^+^ TET2/DNMT3/BRAF subclone was associated to patient #6 brain lesions (**Figure 4D, Table S3**).

Interestingly, single nuclei genotyping (Tapestri) of brain PU.1+ nuclei from patients #8 and #3, who also presented with TET2 and BRAF clones detectable in the brain and blood or bone marrow (**Figure 1B**, **Figure 4C and Table S3**), showed that the TET2 and BRAF variants were mutually exclusive at the single nuclei level (**Figure 4E**), indicating that the patients carried independent hematopoietic clones, both able to colonize the brain. Of note, the TET2 clone was also detected in the unaffected frontal cortex of patient 8 (**Table S3**), indicating that the BRAF clone was a better match with the brain lesions than the TET2 clone.

In contrast to the above, the 2 pediatric-onset LCH patients (patients #1 and #2), did not present with detectable CH, and the BRAF variant was not detected in their white blood cells, even after separation into myeloid and lymphoid cells by flow-sorting (**Figure S6A**) and analyzis by ddPCR at a depth >8000x (**Figure 4C F, Figure S6B, Table S3**). Thus, it is possible to hypothesize that the BRAF mutation has occurred in the resident macrophage lineage, consistent with the general pattern of clonal diversification of microglia (see **Figure 4B**) and as observed in a mouse model^25^. It is also possible that BRAF^V600E^ PU.1+ nuclei originated from a bone marrow clone that spontaneously disappeared, although these 2 patients received minimal chemotherapy, late in the course of the disease and never presented with sign of a myeloproliferative disease (**see** Extended Data Patients).

Similarly, neither CH clones nor the BRAF variant were detected in the bone marrow of patient #5, at a sequencing depth of ∼5000x **(Figure 4C, G, Figure S6B and Table S3**). This patient presented with multifocal histiocytosis in the absence of sign of a bone marrow myeloproliferative disease and did not receive chemotherapy, suggesting again that either that a BRAF^V600E^ bone marrow clone had spontaneously disappeared, or that the BRAF mutation occurred in the resident macrophage lineage.

Patient #7 presented with multiple cancers, and clonal hematopoiesis (TET2 mutation at AF 35%), however, the TET2 variant was undetectable in the brain, while the BRAF variant was undetectable in the bone marrow at a depth of ∼5000x (**Figure 4C and Table S3**), indicating that the brain BRAF clone was independent of his clonal hematopoiesis. A similar pattern was observed in patient #4 (**Figure 4C and Table S3**). These data suggested that, as for patients #1, #2, and #5, the BRAF mutation may have occurred in the resident macrophage lineage **(Figure 4G)**, although it is also possible in theory that a small BRAF^V600E^ bone marrow clone (as observed in patient #3 for example) had spontaneously disappeared in these patients.

These results therefore identified 3 possible different natural histories of the microglial BRAF^V600E^ clones in histiocytosis patients. The brain BRAF clone was a brain subclone of a myeloproliferative disease in one patient, however independent BRAF and TET2 clones can also coexist, at high allelic frequency in the bone marrow and brain. Finally, in the 2 pediatric-onset LCH patients and 3 late onset LCH/ECD patients, the PU.1+ brain BRAF^V600E^ clones may originate from the resident macrophage lineage, although we cannot eliminate the possibility that they originated from transient bone marrow BRAF^V600E^ clones. It is also of note clinical symptoms and the topography of the disease were independent of the putative origin of the BRAF mutant clones.

### Conserved BRAF^V600E^ -driven inflammatory microglia in patients and mice

PCA and pathway analysis of differentially expressed genes in RNAseq from FACS-sorted microglia from the cortex and brainstem of experimental and control mice confirmed that, in addition to the proliferative signature, microglia from the brainstem of mutant mice also presented with strong inflammatory signatures (**Figure 3D, Figure S4 and Table S5**), reminiscent of the transcriptional profile of the patients’ hindbrain (**see Figure 2D**). Accordingly, analysis of common DEG between patients and controls hindbrain and between microglia from the brainstem of experimental and control mice showed a core phagocytic and inflammatory BRAF-associated microglia (BAM) signature characterized by IL1b, NADPH oxidase (Cybb), complement, and phagocytic receptors (**Figure 5A**, **Figure S7A and Table S6**). A single nucleus (sn)-RNAseq analysis of dissected cortex and brainstem from experimental and control mice (**Figure 5B and Figure S7B-E**), confirmed the presence of activated microglia in the brainstem of mutant mice, expressing cathepsins, lysozyme, and APOE (**Figure S7F-I and Table S7**). Overall, these results strongly support the hypothesis that neuro-histiocytosis is a clonal neuro-inflammatory microglial disease associated with the BRAF^V600E^ mutation.

**Figure 5.**
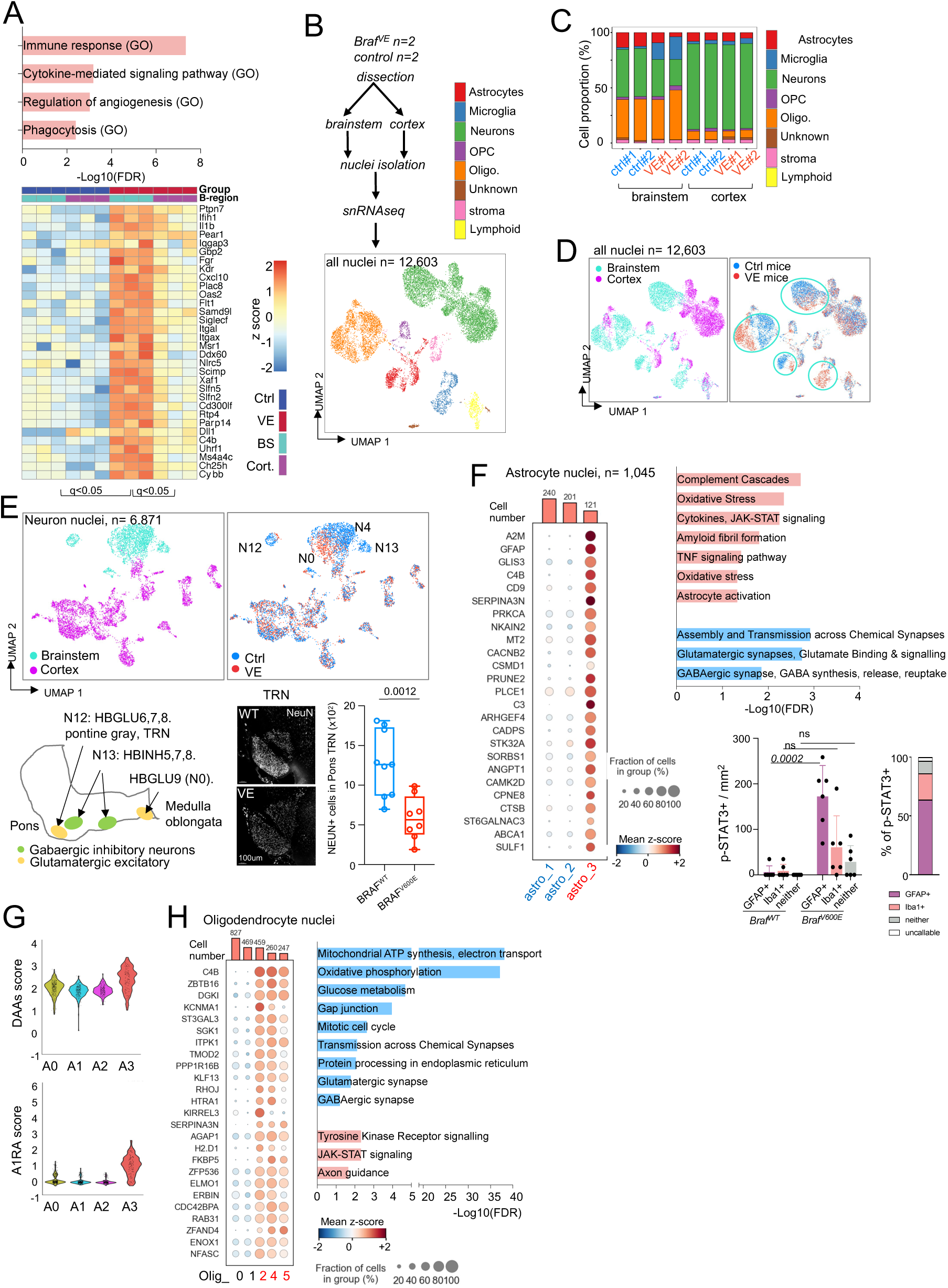
Inflammatory signatures and neuron loss in mouse models of histiocytosis. **(A)** Top, pathway analysis using g:profiler webtool of common differential expressed genes between 2 month old *Cx3cr1^CreERt2^* Braf*^LSL-V600E^* mice microglia in Figure 3 and human whole brain samples in Figure 2. Bottom, hierarchical clustering of common genes in end stage mouse microglia. Expression values are *Z* score transformed. Samples were clustered using average linkage and cluster similarity was determined using the Euclidean distance**. (B)** Single-nuclei RNAseq (snRNAseq) analysis of dissected cortex and brainstem from *Braf^VE/WT^ Cx3cr1^CreER^* mice pulsed with OH-TAM at E9.5 and analyzed at 6 month of age (end stage, VE) (n=2) and littermate controls (Ctrl, n=2). After QC and data processing nuclei were clustered and annotated using Uniform Manifold Approximation and Projection for Dimension Reduction (UMAP) by cell-type. **(C)** Bar plot showing the relative frequency of neurons, microglia and stromal cells by brain region and condition. **(D)** UMAP of all nuclei color-coded by brain region (left) or by condition (right). **(E)** Top, UMAP of neuronal nuclei. color-coded by brain region (left) or by condition (right). Bottom, schematic of brainstem depicting the localization of neuronal clusters reduced in VE samples and NeuN staining (iDISCO) of the pons tegmental nucleus (TRN) from mutants and littermate control. Plot shows the quantification of NeuN staining by immunofluorescence in BRAFV600E and control mice. Each dot represents the mean of three fields per mouse. Statistics: p-values are calculated with Student t test. **(F)** Left, Dot plot showing the expression level (color scale) and the percent of cells expressing (dot size) the most significantly upregulated DEG (log2FC >= 0.5 & FDR <= 0.05) between brainstem astrocytes, A3 (VE) vs A1,2 (control). Barplot represents number of cells per cluster. Right, pathway analysis of DEG in astrocytes using enrichR of upregulated (red) and downregulated (blue) genes in cluster A3 (FDR <0.05, log2FC >= or <= 0.5/-0.5) in comparison to clusters 1,2. Plot shows the quantification of the % of pSTAT3 expressing cells by immunofluorescence, among IBA1+, GFAP+, and IBA1+ GFAP+ cells in the brainstem of *Cx3cr1^CreERT2^; Braf ^LSL-V600E^* and control mice. **(G)** Violin plots showing expression scores for previously defined disease-associated astrocytes signatures, across astrocytes clusters. (**H)** Left, Dot plot showing the expression level (color scale) and the percent of cells expressing (dot size) the most significantly upregulated genes (log2FC >= 0.5 & FDR <= 0.05) between brainstem oligodendrocytes O2,4,5 (VE) vs O 0,1 (control). Right, pathway analysis of DEG in oligodendrocytes using enrichR of upregulated (red) and downregulated (blue) genes in cluster O2,4,5 (FDR <0.05, log2FC >= or <= 0.5/-0.5) in comparison to clusters 0,1.

### BRAF^V600E^ microglia causes massive loss of grey nuclei glutamatergic and GABAergic neurons in the brainstem

(Sn)-RNAseq confirmed that the mutant microglial compartment was expanded at the expense of the neuronal compartment, selectively in the mouse brainstem (**Figure 5C**), reminiscent of changes observed in patients. In addition, nuclei corresponding to all brain cell types from mutant and control mice formed common clusters in the cortex, but distinct clusters in the brainstem, suggesting that all cell types are affected in the brainstem of mutant mice (**Figure 5D**). Analysis of neuronal clusters suggested preferential reduction in numbers of glutamatergic excitatory neurons (HB GLU6,7,8,9, clusters N12 and N0), and GABAergic inhibitory neurons (HBINH5,7,8, cluster N13) from the pons and medulla oblongata grey nuclei, in mutant compared with WT mice (**Figure 5E and Figure S8**). Immunofluorescence analysis of NeuN+ neurons in the pons tegmental reticular nucleus (TRN) confirmed the loss of ∼60% neurons in the TRN of mutant mice (**Figure 5E**). These data confirmed the observation in patients (see **Figure 2**) and allow to conclude that BRAF^V600E^ microglia causes neuronal death in the hindbrain, and specifically the loss of activating and inhibitory neurons within the pons grey nuclei.

### Neurotoxic astrocyte response and reduced oligodendrocyte metabolism

(Sn)-RNAseq analysis also characterized activation of astrocytes from in the brainstem of mutant mice (Cluster Astro_3, **Figure S9 and Table S8)**. This cluster presented with high GFAP expression (**Figure 5F)**, as observed in patients, associated with complement activation, oxidative stress, and JAK-STAT signaling (**Figure 5F-G and Table S8**). Activation of JAK-STAT signaling is reported to promote neuroinflammation in neurodegenerative diseases ^35^, and was confirmed by immunofluorescence (**Figure 5F**). Of note, pathways associated with glutamatergic and GABAergic synaptic processes were downregulated in nuclei from the Cluster Astro_3 (**Figure 5F**), likely in in relation to the decrease in corresponding neurons.

Finally, analysis of oligodendrocytes indicated a global downregulation of metabolism in the brainstem from mutant mice (Oligo clusters O2, O4, O5, **Figure 5H and Table S9**), together with high expression of C4b and α1-antichymotrypsin/Serpina3n (**Figure 5H and Figure S9**), resembling a reactive oligodendrocyte signature previously described as disease associated 1 (DA1) oligodendrocytes in PS2APP and TauPS2APP mice^36^ (**Figure S9**). As observed in astrocytes, pathways associated with glutamatergic and GABAergic synaptic processes were downregulated (**Figure 5H**).

Altogether, these results strongly suggest that mutant microglia promote an astrocytic neurotoxic response which may contribute to neuronal death and predominates in the brainstem grey nuclei and cerebellum in patients and mice. Of note the microglia-driven neurotoxic astrocyte response, described in several human neurodegenerative diseases, and characterized by activation of the JAK-STAT pathway may represent a novel therapeutic target.

### CSF1R inhibition depletes mutant microglia limits neuronal death and improves symptoms and survival

Progressive accumulation of microglial BRAF^V600E^ clones cause neuronal damage and clinical symptoms. Previous studies have shown that BRAF or MEK inhibitors are an effective treatment of LCH and ECD and may delay the onset of neuro-histiocytosis in a mouse model ^25^. However, their effect is only suspensive and long-term treatment carry significant risks of toxicity. We reasoned that if BRAF^V600E^ mutant microglia remained sensitive to depletion by a CSF1R inhibitor, which depletes wild-type microglia *in vivo* ^37^, this treatment may represent an alternative or complement the use of MAP Kinase inhibitors ^38^. We found that treatment with the CSF1R inhibitor PLX5622 ^37^ decreased microgliosis in 6 months old mice more efficiently than treatment with the BRAF Inhibitor PLX4720 ^25,39^ (**Figure 6A**). Treatment with PLX5622 also delayed by several month the onset of neurological symptoms ^25^ (**Figure 6B**), increased survival time (**Figure 6C**), and limited neuronal loss in the TRN (**Figure 6D**), as efficiently as the BRAF inhibitor. Interestingly, treatment with both inhibitors appeared slightly more efficient than each inhibitor separately (**Figure 6B-D)**.

**Figure 6.**
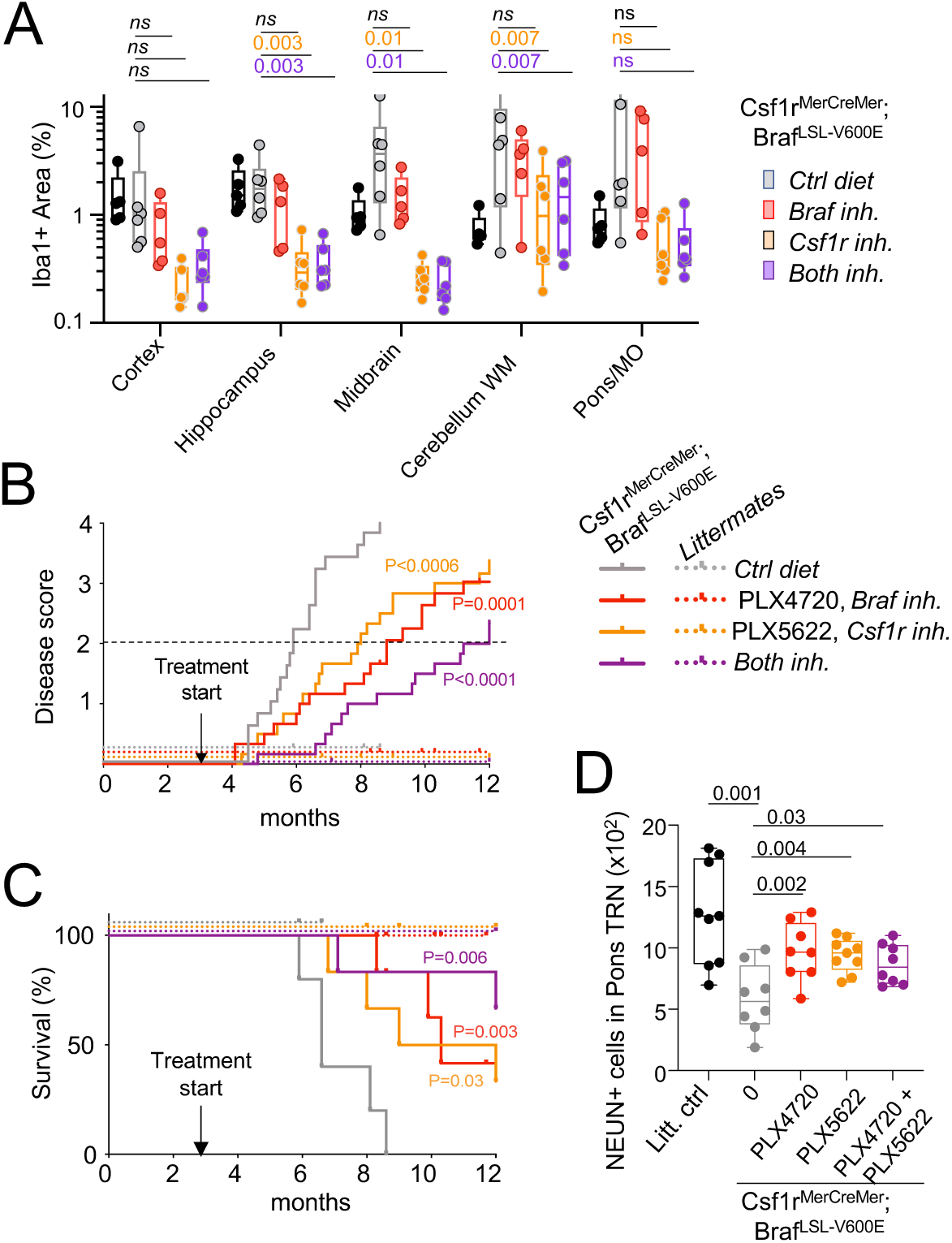
Early microglia depletion with CSF1R inhibitors limits neuronal loss and improves symptoms and survival. **(A)** Percentage of IBA1+ area in brain from *Csf1r^MerCreMer^; Braf^LSL-V600E^*mice and littermate controls pulsed with OH-TAM at E8.5 and treated from 3 months of age with food formulated with CSF1R inhibitor (PLX-5622) n=6, Braf-V600E inhibitor (PLX-4720) n=6, both n=6, or control diet n=5. p-values were calculated with one-way ANOVA, using Dunnett’s multiple comparisons test. **(B)** Disease score progression and **(C)** survival curves for mice in A. Ticks indicate animal death/experimental endpoint. Statistics: Mantel-Cox test. P values are for comparison with control diet. Hazard ratio (logrank): for (a) [CSF1R inh/Ctrl Diet (HR 0.38; 95% CI, 0.19-0.76)], [Braf inh/Ctrl Diet (HR 0.30; 95% CI, 0.15-0.63)], [Both inh/Ctrl Diet (HR 0.20; 95% CI, 0.09 to 0.45)], for (b) [CSF1R inh/Ctrl Diet (HR 0.28; 95% CI, 0.06 to 1.24)], [Braf inh/Ctrl Diet (HR 0.18; 95% CI, 0.03 to 0.92)], [Both inh/Ctrl Diet (HR 0.16; 95% CI, 0.03 to 0.81. **(D)** Quantification of NeuN staining by immunofluorescence in the pons tegmental reticular nuclei (TRN) from control and Csf1r^MerCreMer^; Braf^LSL-V600E^ mice treated with control diet, BRAF inhibitor (PLX4720) CSF1R inhibitor (PLX5622) or the combination. Each dot represents the mean of three fields per mouse. p-values were calculated with ANOVA.

These results suggest that CSF1R inhibitors may represent an alternative to MAP-Kinase inhibitors to limit or prevent neurodegeneration associated with BRAF^V600E^ microglial clones and that the effects of treatment with both BRAF and CSF1R inhibitors may even be additive in terms of survival and disease score, although further studies would be needed to confirm that point.

## Discussion

We report here comprehensive and systematic molecular, pathological, mechanistic and causation studies of the presence and role of mutant microglia in a series of 8 patients diagnosed with BRAF^V600E^ associated pediatric onset LCH and adult onset ECD ^5^. LCH and ECD are rare orphan diseases, justifying the relatively small size of this series of patients. These studies clearly demonstrate that microglia activated by a pathogenic mutation can be neurotoxic in human, an important question which has been recently debated ^40^, and therefore characterize the concept of a neurodegenerative disease mediated by clonal proliferation of inflammatory microglia (CPIM), that might apply to other patients with NDD of unknown mechanisms^29^.

A genetic bar-coding analysis suggests that the cells responsible for the brain lesions can originate from the bone marrow ^18–21^, as demonstrated in 3 patients, but may also arise from the resident macrophage lineage^30,31,41^ as suggested by the analysis of the remaining 5 patients.

Of high potential relevance, these studies identify a neurodegenerative process initiated by pervasive BRAF^V600E^ microglial clones which expand predominantly in the hippocampus, brainstem, and cerebellum, likely due to a local proliferative advantage of mutant BRAF^V600E^ microglia, and irrespective of the putative cellular origin of the microglial clones, and which cause reactive ‘neurotoxic’ astrocytosis and neuronal loss, years before the onset of neurological symptoms. Indeed, the presence of clinical symptoms and extent of neuropathology were correlated with the size of microglia clones, suggesting that incipient neurodegeneration might be targeted by early therapeutic interventions, to prevent or limit the development of a potentially lethal neurodegenerative disease. It is conceivable that advanced imaging protocols, or biomarkers will help to select at risk patients.

Our studies also identify potential molecular targets, as transcriptomics studies identify a BRAF^V600E^ microglia inflammatory and neurotoxic profile ^42–44^, dominated by IL1b, CYBB, complement and phagocytosis, without a clear senescence signature^45–49^, and a neurotoxic astrocyte response reminiscent of other human neurodegenerative diseases, characterized by activation of the JAK-STAT pathway, TNF signaling, oxidative stress and complement production. Finally, we show here a proof of principle that BRAF^V600E^ microglia remain CSF1- dependent, and that microglia depletion with a CSF1R inhibitor^50^ may represent a treatment to prevent neurodegeneration in patients.

## Acknowledgements

This study was supported by the NIH (MSKCC core grant P30 CA008748, 1R01NS115715-01, 1 R01 HL138090-01, 1 R01 AI130345-01 to FG) and the association “Histiocytose France” (www.histiocytose.org). This study was also supported by a Basic and Translational Immunology Grant from the Ludwig Center for Cancer Immunotherapy to FG, and a Cycle for Survival Grant to FG and ELD, RV was supported by the 2018 AACR-Bristol-Myers Squibb Fellowship for Young Investigators in Translational Immuno-oncology 18-40-15-VICA, MP was supported by a post-doctoral fellowship from the Simons society of fellows, EL-R was supported by a Parker Institute for Cancer Immunotherapy Career Development Award. The authors also thank the patients and their families for their agreement to participate to our study, and Anne Schaefer and Pinar Ayata for expert advice on mouse brain anatomy. Sequencing costs of controls individuals were covered in part by a SRA between Neuro-Inflammation NewCo and MSKCC. PLX4720 (BRAF inhibitor), and PLX5622 (CSF1R inhibitor) were kindly provided by Plexxikon Inc). The funders had no role in study design, data collection and analysis, decision to publish, or preparation of the manuscript. This study is dedicated to Florent, a young patient whose years of suffering strengthened our resolve to understand neuro-histiocytosis.

## Authors contribution

FG, RV, JD, designed the study and wrote the draft manuscript. JH, JD, VS, SH, AH, ELD, VT, FC-A, MM, FN-K, AI, NS, ZA, IH, CID took care of the patients, analyzed clinical, biological, and radiological data, and provided biological samples. DS, IP and MR performed neuropathological analysis of patients. RV designed, performed, and analyzed molecular analysis of patients and controls with help from AA AB and OA, SF, HZ, EK, PS, AJ, NDS, AV. MP and EL-R designed, performed, and analyzed murine experiments and snRNAseq with help from CM, YH, SF, and OE. JLC, BB, MO OA-W contributed to data analysis and interpretation and edited the manuscript. All authors contributed to the manuscript.

## Patients and Methods

The study was conducted according to the Declaration of Helsinki, and human tissues were obtained with patient-informed consent and used under approval by the Institutional Review Boards from Memorial Sloan Kettering Cancer Center (IRB protocols #X19-027). Samples from histiocytosis patients were collected under GENE HISTIO study (approved by CNIL and CPP Ile-de France) from Pitié-Salpêtrière Hospital and Hospital Trousseau and from Memorial Sloan Kettering Cancer Center (IRB #14-201). Control frozen brain samples and matching-blood samples were provided by the Netherlands Brain Bank (NBB), the Human Brain Collection Core (HBCC, NIH), the Hospital Sant Joan de Déu, Pitié-Salpêtrière Hospital and the MSKCC Rapid Autopsy Program (IRB #15-021). Samples obtained were clinically and neuropathologically classified by the collaborating institutions as Histiocytoses, and unaffected controls.

**Patients extended description:**

**Patient #1** was a 21-year-old male, diagnosed with LCH in his first year of life after a biopsy of the scalp for a persistent rash -positive for the BRAF^V600E^ mutation-associated with lytic bone lesions of the skull and shoulders on X-rays. He was treated with chemotherapy with good response on the scalp. The patient presented with diabetes insipidus at age 9, at which time a PET scan showed hypometabolism in the cerebellum and the thalamic region^51^ (see **Figure S2B**). At age 13 he was diagnosed with cognitive difficulties, disinhibited behaviour and a cerebellar syndrome and was placed in a special need education programme. MRI at that time shows moderate damage to the pons, dentate nuclei, and the cerebellum (see **Figure S2B**). Insidious progression of cerebellar and cognitive symptoms and new onset motor deficits led him to be wheelchair-bound by the age of 17. At age 18, the patient was briefly treated with the BRAF inhibitor at a dose of 17 mg/kg p.o daily for a week, then at 9 mg/kg for another week before being discontinued for fever hypotonia and pain. The patient did not receive further chemotherapy and died aged 21.

**Neuropathological examination** of the patient brain indicated focal, non-systematized, areas of microglial activation, astrogliosis, and axonal spheroids in the white matter and neuronal loss in grey matter. The lesions predominate in cerebellum, where Purkinje cells are replaced by hyperplasic Bergmann glia, the dentate nucleus, the striatum, the hippocampus (CA4/3) and subiculum, the amygdala and the brainstem, while the frontal and temporal cortex is comparable to control. Histological lesions corresponded to the detection of BRAF^V600E^ in matching brain samples (see **Figure 2)** at 5% to 20% AF indicating that 10% up to 40% of PU.1^+^ cells are mutated in these areas.

**Patient #2** is a 26-year-old female diagnosed with LCH as an infant on a biopsy of persistent scalp rash with no molecular diagnosis. She then progressed with multifocal bone lesions, in zygomatic bone and vertebra, and developed diabetes insipidus. She remained with no evidence of active disease during her childhood and adolescence until the age of 16, when she developed progressively clumsy gait that evolved to cerebellar syndrome with spasticity, and cognitive impairment. With a presumptive diagnosis of neurodegeneration related to LCH she was treated with chemotherapy in her 20’s, followed by IVIG, without clear benefit. MRI at age 26 showed cerebellar lesions similar to patient #1.

**Neuropathological examination:** A core cerebellar biopsy for diagnostic purpose demonstrated regions of Purkinje cell and granule cell loss, microcalcifications and gliosis of the cerebellar white matter. Molecular analysis indicated the presence of BRAF^V600E^ ARAF and KRAS mutations in PU.1+ nuclei at an allelic frequency of 2.5%, indicating 5% mutant cells. Analysis by deep sequencing and ddPCR of the original skin biopsy showed the BRAF^V600E^ mutations at 6% AF, but the ARAF and KRAS mutations were not detectable. Following the cerebellar biopsy and blood samples, the patient was started on treatment with a MEK inhibitor.

**Patient #3** was a 78-old male, born in 1941, and diagnosed with LCH aged 52 in 1993 on a scalp eruption. The patient developed diabetes insipidus at age 59, and later xanthelasmas, bone, cardiac and lung lesions positive for the BRAF^V600E^ mutation, and accordingly diagnosed with ECD. He then developed progressive severe static and kinetic cerebellar syndrome documented at age 77 in 2017. The MRI at the time showed superior cerebellar peduncles abnormalities (**Figure S1E**). The patient then developed abnormal behaviour, and episodes of confusion and falls, and died age 79 of an ischemic and hemorrhagic stroke.

**Neuropathological examination:** Ischemic and hemorrhagic lesions made the neuropathological analysis difficult, but focal, non-systematized areas of microglial activation, astrogliosis, and axonal spheroids and neuronal loss are identified in the cerebellum (dentate nuclei). Molecular analysis indicates the presence of large PU.1^+^ BRAF^V600E^ clones (AF: 10 to 20%) in the cerebellum, brainstem and hippocampus, similar to patient #1 and smaller clones (AF: ∼1% in the frontal and temporal cortex. Taken together, neurological, histological and molecular analysis of these 3 patients suggest that the cerebellar, pyramidal and cognitive syndrome are associated with histological lesions involving primarily the cerebellum, brainstem and hippocampus, and large PU.1^+^ BRAF^V600E^ clones in 5 to 40% brain macrophages in these structures.

**Patient #4** was a 66-year-old male diagnosed with ECD with a BRAF^V600E^ mutation in the skin at age 59, with orbital lesions, multifocal bone disease, massive retroperitoneal fibrosis, along with lung and cardiac involvements. He was treated initially with IFNa and subsequently with the BRAF inhibitor vemurafenib. He was diagnosed with pancreatic cancer at age 66 and died the same year. The patient did not present any neurological symptoms, and specifically did not have any cerebellar signs nor cognitive complain, however an MRI performed in 2016 showed hypersignal in the dentate nucleus.

**Neuropathological analysis** revealed severe focal, non-systematized, areas of microglial activation, astrogliosis, axonal spheroids and neuronal loss in the pons, and milder lesions in hippocampus and cerebellum, which correspond to the detection of a large PU.1^+^ BRAF^V600E^ clones in the pons (AF ∼10%), a smaller clone in the hippocampus (AF ∼1%) and no detectable clone in the cerebellum. The BRAF^V600E^ clone was also undetectable in 2 distinct bone marrow samples, and conversely a bone marrow DNMT3 mutant clone was not detectable in the brain.

**Patient #5** was a 78-year-old female diagnosed with LCH at age 75 on a skin eruption. She had skin, bone, lung and cardiac involvements and a severe sclerosing cholangitis with a BRAF^V600E^ mutation identified in a liver biopsy, but no neurological disease, and died of hepatic insufficiency. The patient did not receive chemotherapy because of her hepatic insufficiency. MRI performed at diagnosis showed bilateral dentate nuclei T2 hypersignal **(Figure S2I)**.

**Neuropathological analysis** revealed severe histological lesions in the dentate nuclei, hippocampus (junction CA1/2), pons and medulla oblongata. Molecular analysis showed corresponding large (AF∼10%) PU.1^+^ BRAF^V600E^ clones in the hippocampus, midbrain, pons and medulla oblongata, and smaller (1%) PU.1^+^ BRAF^V600E^ clones in the cerebellum and temporal lobe **(Figure 2).**

**Patient #6** was a 61-year-old male diagnosed with ECD at age 59, based on skin and pulmonary involvement and hypermetabolic bone lesions. Skin biopsy confirmed a BRAF^V600E^ mutation. The patient was also diagnosed with clonal hematopoiesis (DNMT3A, JAK2, TET2 at ∼50% AF). and presented a splenic infarction. Due to the multisystem character of his histiocytic disease, he was given 4 months the MEK inhibitor cobimetinib in early 2019, but developed a T-cell lymphoma, responsible for his death age 61. A distal motor deficit graded MRC (Medical Research Council) 4/5 was detected on lower limbs with abolition of deep tendon reflexes. Electromyography ruled out a multiple mononeuropathy and nerve compression. No signs of central nervous system involvement including cerebellar and cognitive involvement was observed. MRI at age 60 also showed small and multiple dentate nuclei T1 hyperintensities.

**Neuropathological analysis** found focal, non-systematized, areas of microglial activation, and astrogliosis in the pons and cerebellum. Molecular analysis indicated the presence of 2 small PU.1^+^ BRAF^V600E^ clones in the cerebellum and pons. The bone marrow clone carrying DNMT3A/ JAK2/ TET2 mutations at AF ∼50% was also detected in the brain, at AF 2% to 9% (average 7%).

**Patient #7** was a male with a history of kidney adenocarcinoma and prostatic carcinoma, diagnosed with ECD at age 85 on a biopsy of a retroperitoneal fibrosis. The patient also presented with hypermetabolic long bone lesions. The patient developed lung metastasis of the kidney adenocarcinoma. Brain CT scan diagnosed an acute left hemispheric subdural hematoma after a fall age 88 followed by subacute confusion. A significant neurological improvement was observed after neurosurgical removal of the hematoma. Follow-up brain CT showed chronic subdural hematoma that did not require surgical procedure. Persistent memory disturbances and balance disorders were reported afterwards and attributed to the subdural hematoma. The patient also carried a clonal hematopoiesis with a TET2+ clone in 34% of BM cells. The patient died of denutrition and nosocomial infection at age 89.

**Neuropathological analysis:** found focal, non-systematized, areas of microglial activation, and astrogliosis in the pons and cerebellum Molecular analysis indicated the presence of a PU.1^+^ BRAF^V600E^ clone in the pons **(Figure 2)**. Clonal hematopoiesis was established on a TET2 variant at AF ∼35% in the bone marrow, but the clone was not detected in the brain.

**Patient #8** was a female with an history of bilateral metastatic breast cancer first diagnosed at age 48, treated with tumorectomy, bilateral mastectomy, irradiation and chemotherapy. She was subsequently diagnosed with ECD 10 years later with skin, bone, heart and retroperitoneal lesions, diabetes insipidus. She was treated with the BRAF inhibitor vemurafenib with a durable an sustained reponse ^52^. The patient also presented with a myeloproliferative syndrome (TET2+, AF 30%) **(Figure 1)**. A static and kinetic cerebellar syndrome and cognitive disturbances with dysexecutive syndrome developed at age of 66. MRI documented brain lesions predominant in the cerebellum. The patient died of infection in the context of metastatic breast cancer and neurodegeneration at age 68.

**Neuropathological analysis:** found focal, non-systematized, areas of microglial activation, and astrogliosis in all brain regions examined except the frontal cortex. Molecular analysis indicated the presence of PU.1^+^ BRAF^V600E^ clones in the the same regions. Clonal hematopoiesis was established on a TET2 variant at AF ∼30% in the bone marrowand was also detected in the brain, albeit at lower AF.

## Methods

### Nuclei isolation from frozen brain samples

All samples were handled and processed under Air Clean PCR Workstation. ∼250-400 mg of frozen brain tissue were homogenized with a sterile Dounce tissue grinder using a sterile non-ionic surfactant-based buffer to isolate cell nuclei (250 mM Sucrose, 25 mM KCL, 5 mM MgCl2, 10 mM Tris buffer pH 8.0, 0.1% (v/v) Triton X-100, 3 μM DAPI, Nuclease Free Water). Homogenate was filtered in a 40-μm cell strainer and centrifuged 800g 8 min 4°C. To clean-up the homogenate, we performed a iodixanol density gradient centrifugation as follow: pellet was gently mixed 1:1 with iodixanol medium at 50% (50% Iodixanol, 250 mM Sucrose, 150 mM KCL, 30 mM MgCl2, 60 mM Tris buffer pH 8.0, Nuclease Free Water) and homogenization buffer. This solution layered to a new tube containing equal volume of iodixanol medium at 29% and centrifuged 13.500g for 20 min at 4°C. Nuclei pellet was gently resuspended in 200 μl of FACS buffer (0.5% BSA, 2mM EDTA) and incubated on ice for 10 min. After centrifugation 800g 5 min 4°C, sample was incubated with anti-NeuN (neuronal marker, 1:500, Anti-NeuN-PE, clone A60 Milli-Mark™) for 40 min. After centrifugation 800g 5 min 4°C, sample was washed with 1X Permeabilization buffer (Foxp3 / Transcription Factor Staining Buffer Set, eBioscience™) and centrifuged 1300g 5 min, without breaks to improve nuclei recovery. Staining with anti-Pu.1 antibody in 1X Permeabilization buffer (microglia marker 1:50, PU.1-AlexaFluor 647, 9G7 Cell Signaling) was performed for 40 min. After a wash with FACS buffer sample were ready for sorting. Nuclei are FACS-sorted in a BD FACS Aria with a 100-μm nozzle and a sheath pressure 20 psi, operating at ∼1000 events per second Pellet is gently resuspended in 200 μl of FACS buffer (0.5% BSA, 2mM EDTA) and incubated on ice for 10 min. After centrifugation 800g 5 min 4°C, sample was incubated with anti-NeuN (neuronal marker, 1:500, Anti-NeuN-PE, Milli-Mark™) for 40 min. After centrifugation 800g 5 min 4°C, sample is washed with 1X Permeabilization buffer (Foxp3 / Transcription Factor Staining Buffer Set, eBioscience™) and centrifuged 1300g 5 min, without breaks. Staining with anti-Pu.1 antibody in 1X Permeabilization buffer (microglia marker 1:50, PU.1-AlexaFluor 647, 9G7 Cell Signaling) was performed for 40 min. After a wash with FACS buffer sample were ready for sorting. Nuclei are FACS-sorted in a BD FACS Aria with a 100-μm nozzle and a sheath pressure 20 psi, operating at ∼1000 events per second. For purity analysis, snRNAseq was performed on sorted PU.1+ nuclei from one control brain, with a resulting purity of >93% microglia ^29^. For DNA sequencing, nuclei were sorted into 1.5 ml certified RNAse, DNAse DNA, ATP and Endotoxin free tubes containing 100 μl of sterile PBS. For each population we sorted >10^5^ nuclei. Nuclei pellets were centrifuged 20 min at 6000g and processed immediately for gDNA extraction with QIAamp DNA Micro Kit (Qiagen) following manufacture instructions. DNA from whole-blood or bone marrow samples was extracted using the same protocol. Flow cytometry data was collected using DiVa 8.0.1 Software. Subsequent analysis was performed with FlowJo_10.6.2.

### Library preparation and sequencing

DNA samples were submitted to the Integrated Genomics Operation (IGO) at MSKCC for quality and quantity analysis, library preparation and sequencing. After PicoGreen quantification, ∼200ng of genomic DNA were used for library construction using the KAPA Hyper Prep Kit (Kapa Biosystems KK8504) with 8 cycles of PCR. After sample barcoding, 2.5ng-1µg of each library were pooled and captured by hybridization with baits specific to either the HemePACT (Integrated Mutation Profiling of Actionable Cancer Targets related to Hematological Malignancies) assay, designed to capture all protein-coding exons and select introns of 576 (2.88Mb) commonly implicated oncogenes, tumor suppressor genes and or HemeBrainPACT (716 genes, 3.44 Mb) an expanded panel that included additional custom targets related to neurological diseases (see Supplementary Table 2).Capture pools were sequenced on the HiSeq 4000, using the HiSeq 3000/4000 SBS Kit (Illumina) for PE100 reads. Samples were sequenced to a mean depth of coverage of 1106x (Control samples: 1088Xx, Histiocytosis samples 1178x).

### Mutation data analysis

The data processing pipeline for detecting variants in Illumina HiSeq data is as follows. First the FASTQ files are processed to remove any adapter sequences at the end of the reads using cutadapt (v1.6). The files are then mapped using the BWA mapper (bwa mem v0.7.12). After mapping the SAM files are sorted and read group tags are added using the PICARD tools. After sorting in coordinate order the BAM’s are processed with PICARD MarkDuplicates. The marked BAM files are then processed using the GATK toolkit (v 3.2) according the best practices for tumor normal pairs. They are first realigned using ABRA (v 0.92) and then the base quality values are recalibrated with the BaseQRecalibrator. Somatic variants are then called in the processed BAMs using MuTect (v1.1.7) for SNV and ShearwaterML ^53,54^.

### MuTect1

to identify somatic variants and eliminate germline variants, we run the pipeline as follow: PU.1, DN and Blood samples against matching-NeuN samples, and NeuN samples against matching-PU.1. In addition, we ran all samples against a Frozen pool of 10 random genomes. We selected Single Nucleotide Variations (SNVs) mutations [Missense, Nonsense, Splice Site, Splice Regions] that were supported by at least 4 or more mutant reads and with coverage of 50x or more. Fill-out file for each project (∼27 samples per sequencing pool), allowed to exclude mutations with high background noise. This resulted in 612 mutations (missense, nonsense, splice_site, splice_region).

**ShearwaterML**, was used to look for low allelic frequency somatic mutations as it has been shown to efficiently call mutations present in a small fraction of cells with true positives being ∼90%. Briefly, the basis of this algorithm is that is uses a collection of deep-sequenced samples to learn for each site a base-specific error model, by fitting a beta-binomial distribution to each site combining the error rates across all normal samples both the mean error rate at the site and the variation across samples, and comparing the observed mutation rate in the sample of interest against this background model using a likelihood-ratio test. For detailed description of this algorithm please refer to^53,54^. In our data set, for each cell type (NeuN, DN, PU.1) we used as “normal” a combination of the other cell types (from Histiocytosis as well as control samples), i.e PU.1 vs NeuN+DN, DN vs NeuN+PU.1, NEUN vs PU.1+DN, Blood vs NeuN+DN. Since all samples were processed and sequence the same way, we expect the background error to be even across samples. More than 400 samples were used as background leading to an average background coverage >400.000x. Resulting variants for each cell type were filtered out as germline if they were present in more than 20% of all reads across samples. Additionally, mutations with coverage of less than 50x and more than 35% variant allelic frequency (VAF) were removed from downstream analysis. p-values were corrected for multiple testing using Benjamini & Hochberg’s False Discovery Rate (FDR) ^55^ and a q-value of cutoff of 0.01 was used to call somatic mutations. Mutations were required to have a least one supporting read in each strand. Somatic mutations within 10bp of an indel were filtered out as they typically reflect mapping errors. We selected Single Nucleotide Variations (SNVs) mutations [Intronic, Intergenic, Missense, Nonsense, Splice Site, Splice Regions] that were supported by at least 4 or more mutant reads and annotated them using VEP. Finally, we excluded mutations with a MAF (minor allelic frequency) cutoff of 0.01 using the gnomeAD database. This resulted in 424 SNVs.

We compared the final mutant calls of MuTect and ShearwaterML and found that >20% of the events (137 mutations) called by MuTect1 were also called by ShearwaterML. The aggregation of MuTect and ShearwaterML resulted in a total of 899 unique variants, with a mean coverage at the mutant site of 659x (10% percentile: 313x, 90% percentile: 1095x) and a mean of 20 mutant reads (10% percentile: 4, 90% percentile: 42), and 87% of mutated supported by at least 5 mutant reads (mean variant allelic frequency of 3%).

### Validation of mutations

We performed validation of 7.56% of mutations (82/899) by droplet-digital-PCR (ddPCR) on pre-amplified DNA or on libraries (when DNA not available) and confirmed 81/82 of mutations tested (>98%). Most assays were performed in all cell types isolated from the same brain region and matching blood/bone marrow). The mean depth of ddPCR was ∼4000x. For BRAFV600E ddPCR we used BRAF_V600E Bio-Rad validated assay (Unique Assay ID: dHsaMDV2010027). Other assays specific for the detection of mutations were designed and ordered through Bio-Rad. For newly designed assays, cycling conditions were tested to ensure optimal annealing/extension temperature as well as optimal separation of positive from empty droplets. All reactions were performed on a QX200 ddPCR system (Bio-Rad catalog # 1864001). When possible, each sample was evaluated in technical duplicates or quartets. Reactions contained 10ng gDNA, primers and probes, and digital PCR Supermix for probes (no dUTP). Reactions were partitioned into a median of ∼31,000 droplets per well using the QX200 droplet generator. Emulsified PCRs were run on a 96-well thermal cycler using cycling conditions identified during the optimization step (95°C 10’; 40-50 cycles of 94°C 30’ and 52-56°C 1’; 98°C 10’; 4°C hold). Plates were read and analyzed with the QuantaSoft sotware to assess the number of droplets positive for mutant DNA, wild-type DNA, both, or neither.

### Quantification of mutational load and statistics

We defined mutational load or mutational burden as the number of synonymous and non-synonymous somatic single-nucleotide-variations (SNV) per megabase of genome examined ^56^. To quantify mutational load we took into consideration the panel used for sequencing each sample: HemePACT (2.88 Mb) or the extended panel HemeBrain-PACT (3.44 Mb) (see Supplementary Table 1-3). Therefore, the number of mutations was normalized by the number of Mb sequenced for that specific sample. In the cases where we calculated mutational load per patient, we averaged the mutational load of each sample from that patient for a given cell type [(i.e if for one patient, 2 PU.1 samples were sequenced, one from hippocampus and one from cortex (with BRAIN-PACT) then the mutational load for PU.1 for that patient is the mean of the mutational load of the 2 PU.1 samples analyzed). Statistical significance was analyzed with GraphPad Prism (v9) and R (3.6.3). Non-parametric tests were used when data did not follow a normal distribution (Normality test: D’Agostino-Pearson and Shapiro-Wilk test). In all the statistical tests, significance was considered at P < 0.05. For Venn Diagram plots, we used ^57^.

### Single-nuclei library preparation and sequencing (Tapestri)

PU.1+ nuclei from Pons and Cerebellum from patient #6, Cerebellum from patient #3 and Pons from patient #8 were isolated as detailed above and processed as in ^58^. Briefly, nuclei were suspended in Mission Bio cell buffer at a maximum concentration of 4000 nuclei/µl, encapsulated in Tapestri microfluidics cartridge, lysed and barcoded. Barcoded samples were then put through targeted PCR amplification with the Myeloid panel (MB03-0036, Mission Bio) which included the mutations to be tested. PCR products were removed from individual droplets, purified with Ampure XP beads and used as templates for PCR to incorporate Illumina i5/i7 indices. PCR products were purified again, quantified with an Agilent Bioanlyzer for quality control, and sequenced on an Illumina NovaSeq. The minimum total read depth was determined by same formula as used in ^58^. A total of 2769 nuclei were analyzed from patient #6, 4919 from patient #3 and 710 from patient #8.

### Demultiplexing and alignment

FASTQ files for single-nuclei DNA libraries were first processed through Mission Bio’s Tapestri pipeline v3 part1 with default parameters for demultiplexing and alignment that resulted in two outputs:

1. Single-barcode BAM files
2. Barcode-by-amplicon read-count matrix *X*1. This matrix is meant for single-cell copy number analysis, which was not relevant for this study.

Briefly, the Mission Bio pipeline v3 part1 proceeds in the following steps:

1. Adaptor sequences are trimmed from the reads; barcode sequences are extracted; the reads are aligned to the hg19 genome (UCSC)
2. The reads are assigned to individual cell barcodes while filtering out the low-quality cell barcodes.
3. For each barcode, the number of forward reads aligned to each amplicon is used to form matrix *X*1.

A more detailed documentation of the Mission Bio pipeline is available at: https://support.missionbio.com/hc/en-us/categories/360002512933-Tapestri-Pipeline and ^59^.

### Single-nuclei library preparation and sequencing (Tapestri)

PU.1+ nuclei were isolated as detailed above and processed as in ^58^. Briefly, nuclei were suspended in Mission Bio cell buffer at a maximum concentration of 4000 nuclei/µl, encapsulated in Tapestri microfluidics cartridge, lysed and barcoded. Barcoded samples were then put through targeted PCR amplification with the Myeloid panel (MB03-0036, Mission Bio) which included the mutations to be tested. PCR products were removed from individual droplets, purified with Ampure XP beads and used as templates for PCR to incorporate Illumina i5/i7 indices. PCR products were purified again, quantified with an Agilent Bioanlyzer for quality control, and sequenced on an Illumina NovaSeq. The minimum total read depth was determined by same formula as used in ^58^.

### Demultiplexing and alignment

FASTQ files for single-nuclei DNA libraries were first processed through Mission Bio’s Tapestri pipeline v3 part1 with default parameters for demultiplexing and alignment that resulted in two outputs:

1. Single-barcode BAM files
2. Barcode-by-amplicon read-count matrix *X*1. This matrix is meant for single-cell copy number analysis, which was not relevant for this study.

Briefly, the Mission Bio pipeline v3 part1 proceeds in the following steps:

1. Adaptor sequences are trimmed from the reads; barcode sequences are extracted; the reads are aligned to the hg19 genome (UCSC).
2. The reads are assigned to individual cell barcodes while filtering out the low-quality cell barcodes.
3. For each barcode, the number of forward reads aligned to each amplicon is used to form matrix *X*1.

A more detailed documentation of the Mission Bio pipeline is available at: https://support.missionbio.com/hc/en-us/categories/360002512933-Tapestri-Pipeline and ^59^.

### Single-cell variant calling and joint genotyping

We used a custom variant calling pipeline [Zhang et al., *bioRxiv* 2024]. Briefly:

1. Single-cell level variant calling for discovery: Mutect2 (GATK 4.2.5.0) and custom filters are used for *de novo* mutation calling and filtering at single-cell level. All cells’ variant lists are merged into a consensus list.
2. Sample-level joint genotyping: bcftools (v1.11) is used for genotyping across all cells in the same sample using the consensus list from above. This results in a second matrix *X*2, which is variant-by-cell.
3. Patient-level joint genotyping: If multiple samples of the same patient are surveyed, variant lists from the samples are merged into a consensus list for genotyping across all cells of that patient.

### Analysis of clonal relationship between a pair of mutations

Given two SNVs’ distribution of reads across single cells, we developed a statistical measure of their clonal relationship – whether they are colocalizing in the same set of cells or mutually exclusive. Suppose we have a probability matrix *P* ɛ [0,1]*^n^*^×*m*^ for *n* cells and *m* mutations, where *p_i,j_* is the probability that mutation *j* is present in cell *i* . We will describe how the probability is modeled in the next section. For any two mutations *j* and *j*’, let *Y* be the random variable that denote the number of observations of mutual exclusivity across the *n* cells, i.e. the number of observations of either (0,1) or (1,0) gametes in the mutation matrix. Assuming that the mutation states of cells are independent, *Y* is a Poisson binomial variable with parameters:

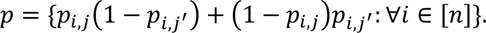

We define a null distribution that conserves the marginal expectation of the number of occurrences of each mutation across the *n* cells.

Specifically, we define probability matrix *P*^’^ such that:

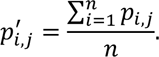

Let *X* be the random variable that denote the number of observations of mutual exclusivity for mutations *j* and *j* ^’^ across the *n* cells under the null distribution.

*X* is a binomial variable with *n* trials and probability of success given by:

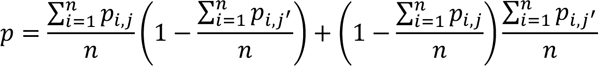

The p-value for rejecting the null hypothesis is given by *Pr*(*X* ≥*Y*). We formally define hypothesis testing problem as follows:

Given a probability matrix *P*and two mutations *j* and *j*^’^, find the p-value Pr(*X* ≥ *Y*), where *Y* and *X* is the number of occurrences of gametes (0,1) and (1,0) for mutation *j* and *j*^’^ under the alternate and the null hypothesis, respectively.

We compute the p-value Pr(*Y* ≤ *X*) by marginalizing *X* as follows,

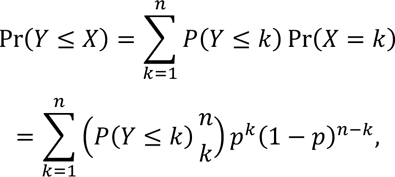

where Pr(*Y* ≤ *k*) is the cumulative probability of the Poisson Binomial variable. If we want to test for occurrence of different gametes, we simply modify the indicators in the model: i.e. if we testing for colocalization, we would set the gametes to be (0,0) and (1,1).

### Read count model

We use the mutation genotype model used by ConDoR to model the probability of a mutation being present in a cell ^60^. We assume that measurement of mutation across cells is independent. As such, let *s_i,l_* denote the state of mutation *j* in copy number cluster *l*; let *q_i,j_* and *r_i,j_* denote the variant reads and total reads of mutation *j* in cell *i*, respectively. We use the beta-binomial model for the read counts *r_i,j_* of each mutation in each cell. When the mutation is present, i.e. *s_i,j_* = 1, we model the read counts as:

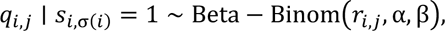

where we set α= β = 1 for this study. When the mutation is absent, i.e. s_i,j_ = 0, there may still be variant reads, i.e. *r_i,j_* > 0, because of sequencing errors. Let *f_p_* be the sequencing error rate to produce a false positive variant read and *d* be the dispersion parameter. We model the read counts for *s_i,j_* = 0 case as follows,

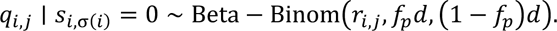

We set *f_p_* = 0.001 and *d* = 15.

### Mixed-effects linear regression model

Age and neurological status were incorporated as fixed effects, while donor as random effects. We also tested sex, brain region, or the combination of sex and brain region as additional random effects, but these variables did not improve the overall model fitting as determined by likelihood ratio test (P > 0.97). In this model, neurological status is a significant predictor of SNV burden (P = 3e-4 by likelihood ratio test). Age is also a predictor (P=0.0035 by likelihood ratio test).

### Neuro-histological analysis of human samples

Neuropathological analysis of human brains was performed by 3 of us (DS, IP, MR). Immunohistochemistry of tissue of patients and controls was carried out on 3–4-μm thick paraffin sections, fixed with 4% formaldehyde. Hematoxylin and eosin and immunohistochemical analysis with rabbit anti-IBA1 (1:500, 019-19741, Wako), GFAP (1:500, 6F2, Dako) was performed on paraffin sections, in Ventana XT platform.

### RNA extraction from human brain samples, library preparation and sequencing

RNA was extracted from frozen tissue using the Qiagen all-prep DNA/RNA mini kit (80204) according to the manufacturer’s instructions. RNAs were used for ribogreen quantification and quality control on Agilent BioAnalyzer. Subsequently, 500 ng of total RNA was used for polyA selection and Truseq library preparation according to the instructions provided by Illumina (TruSeq RNA Sample Prep Kit v.2), with 8 cycles of PCR. Samples were barcoded and run on a Hiseq 4000 in a 125 bp–125 bp paired-end run, using the TruSeq SBS Kit v.3 (Illumina). An average of 75 million paired reads was generated per sample.

### Analysis of human and mice bulk RNAseq

FastQ files of 2x100bp paired-end reads were quality checked using FastQC (https://www.bioinformatics.babraham.ac.uk/projects/fastqc/, 2012). Samples were with high quality reads (Phred score >= 30) were aligned to the Human reference genome (GRCh378/hg38) using STAR aligner (version 2.7) ^61^. Gene quantification was performed using feature counts from the Subread package in R ^62^. Aligned reads were visualized using IGV (version 2.8.9). Gene expression levels were normalized and log2 transformed using the Trimmed Mean of M-values (TMM) method. Differential expression analysis was performed using the edgeR package in R ^63^. Significantly differentially expressed genes were selected with controlled False Positive Rate (B&H method) at 5% (FDR <= 0.05). Upregulated genes were selected at a minimum log2 fold change of 2 and downregulated genes at a minimum log_2_ fold change of -2. Heatmaps were drawn on the normalized expression matrix using the pheatmap package in R. Euclidean distance similarity metric and complete linkage algorithm were used for hierarchical clustering. Gene ontology (GO) and pathway (WikiPathways, KEGG, Reactome) analysis using g:profiler webtool was performed on the list of upregulated genes (log_2_ >= 2 and FDR <=0.05) ordered by importance ^64^. GO terms and pathways were selected based on an FDR <= 0.05.

### Mouse models

*Csf1r^MeriCreMer^* mice ^22^ (FVB/NJ) were kindly provided by Dr Jeffrey Pollard, *Cx3cr1^CreER^*mice ^65^ (C57BL/6J, Jackson Stock No. 021160) (kindly provided by Dr Dan Littman), *Rosa26^LSL-YFP^* mice ^66^, and *Braf ^LSL-V600E^* mice ^66^ (C57BL/6J) were kindly provided by C. Pritchard (Leicester, UK) were previously described, including genotyping protocols. Mice were maintained at Memorial Sloan Kettering Cancer Center (MSKCC) Zuckerman Research Center animal facility under specific-pathogen-free conditions. All experiments were performed according to the guidelines set by the Institutional Animal Care and Use Committee as well as the National Institutes of Health Guide for the Care and Use of Laboratory Animals, Institutional Review Board (IACUC 15-04-006 and 13-04-003) from MSKCC. *Braf ^WT^cre^−^*, *Braf ^V600E^cre^−^* and *Braf ^WT^cre^+^* littermates were considered Braf WT for representation of data. For targeting of BRAF(V600E) mutation to microglia progenitors, we genetically targeted EMPs by pulse-labelling *Csf1r^MeriCreMer^;Braf ^LSL-V600E^;Rosa26^LSL-YFP^* E8.5 embryos as previously described ^22,23^ or *Cx3cr1^CreER^;Braf ^LSL-V600E^*E9.5 embryos (with a single injection of 37.5 mg per kg (body weight) of 4-hydroxytamoxifen (4-OHT, Sigma-Aldrich) into pregnant females. To target of BRAF(V600E) mutation after birth to CX3CR1positive cells *Cx3cr1^CreER^;Braf ^LSL-V600E^* mice were pulsed at 1 month of age with 37.5 mg per kg (body weight) of Tamoxifen (Sigma-Aldrich) intraperitoneally for 5 consecutive days . EMPs appear in the mouse yolk sac at embryonic day E8.5 and express the Csf1 receptor (Csf1r) and one day later (E9.5) express the chemokine receptor CX3CR1^22–24^. They colonize the fetal liver from E9.5 and give rise to macrophage precursors that distribute in embryonic tissues and differentiate into tissue-specific macrophage subsets, such as microglia in the central nervous system ^22^. Embryonic development was estimated considering the day of vaginal plug formation as 0.5 days post-coitum (dpc). A short treatment with 4-OHT leads to transient nuclear translocation of the estrogen receptor-Cre-recombinase fusion protein (MeriCreMer or CreER) in cells expressing the *Csf1r^MeriCreMer^*transgene or *Cx3cr1^CreER^* Knock-in allele and deletion of a floxed stop cassette (LSL) in the *Braf^LSL-V600E^* and *Rosa26^LSL-YFP^* alleles. 4-OHT was supplemented with 18.75 mg per kg (body weight) progesterone (Sigma-Aldrich) to counteract the mixed oestrogen agonist effects of tamoxifen, which can result in fetal abortions.

### Flow cytometry of mouse microglia, cell sorting and RNA isolation for bulk-RNAseq

Animals where lethally anesthetized by an intravenous injection (IV) of 10 μl/g body weight of ketamine, xylazine, acepromazine and followed by a transcardial perfusion with 20 ml ice cold PBS (gibco 14190-144). Brain was dissected and regions of one hemisphere were further separated into the Cortex, Striatum, Hippocampus, Midbrain/Interbrain, Brainstem (Pons/Medulla), Cerebellum and cervical spinal cord and placed on ice in PBS. Brain areas were individually Dounce homogenized in 6ml ice cold FACS buffer (5% Bovine Serum Albumin Proteins, 2 mM Ethylenediaminetetraacetic Acid (EDTA) in PBS (Gibco 14190-144), sterile filtered) 10-20 times with the loose and then the tight pestles. Homogenate was strained through a 100 μm cell strainer (Falcon 352360) into a 15 ml falcon and Douncer was washed out with 2 ml FACS buffer and strained as well into the 15 ml falcon. The single cell suspension was centrifuged at 300 g for 5 min at 4 °C. Resulting cell pellet was resuspended in ice cold PBS-buffered 40% Percoll^TM^ (GE Healthcare 17-0891-01 and centrifuged for 30 min at 500 g at 4 °C with full acceleration and braking. Supernatant was discarded and cell pellet was washed once with 10 ml cold FACs buffer centrifuged again for 5 min at 300 g at 4 °C. An aliquot of the cell pellet was used for RNA isolation with Qiagen all-prep DNA/RNA mini kit (80204) according to the manufacturer’s instructions. Remaining cell pellet was resuspended in 50 μl antibody staining mix and transferred to a 96 well plate to incubate for 30 min on ice.

Antibody staining mix contained FACs buffer with F4/80 (APC) 1:100, CD45 (APC-Cy7) 1:100, CD115 (PE) 1:100, CD11b (PE-Cy7) 1:200, and CD16/32 blocking 1:100 diluted. Samples were spun down at 300 g for 5 min at 4 °C and then resuspended in 200 μl FACs buffer with 0.1 ug/mL DAPI and kept on ice. Stained cells from separated brain areas were individually sorted for CD11b^+^, CD45^int^ microglia with the BD FACSAria II in purity mode using a 100 μm nozzle into individual Eppendorfs filled with 750 μl FACS buffer.

### DNA extraction from mouse microglia

DNA was isolated from sorted microglia by resuspending cell pellet in 10 μl or 20 μl QuickExtract (20 μl for samples with more than 50 000 cells) and pulse-vortexed for 15 seconds. Samples were incubated for 6 min at 65 °C, followed by vortexing for 15 s. Reaction was halted by incubating samples for 2 min at 98 °C. All samples were diluted with 20 μl water and stored at −20 °C until submission to the Integrated Genomics Operation facility of MSKCC for digital PCR.

### Digital PCR for Braf V600E allelic frequency in mice

Recombination frequency of the *Braf^LSL-V600E^* allele was assessed in pooled microglia samples from each brain region using a multiplex digital PCR (ddPCR) assay detecting the novel junction formed after Cre-mediated recombination (Braf-Rec) and the targeted unrecombined allele (Braf-minigene). The endogenous untargeted allele was not detected by the assay. Recombination frequency was calculated as (counts Braf-Rec)/(counts Braf-Rec + counts Braf-minigene). Because experimental mice have one germline copy of *Braf^LSL-V600E^*, mutant microglia frequency was expressed as Allelic Frequency = Recombination frequency/2. To confirm equal efficiency for the two assays, a gBLOCK(IDT) containing the Braf-Rec and Braf-minigene amplicons separated by a KpnI restriction enzyme site was used as a 1:1 positive control. The gBLOCK was digested with KpnI and purified using the Qiagen Reaction Cleanup Kit prior to use. Genomic DNA from WT and *Braf^LSL-V600E^* Cre^-^ mice were run as negative controls.

### Mouse tissue preparation for histology

Dissected brain hemisphere and whole vertebra was fixated for 24 h in 4% Paraformaldehyde in PBS at 4 °C. Tissues were washed for 10 min 3 times in PBS at room temperature. After the fixation, the spinal cord was dissected out of the vertebra. Tissue was stored in 70 % Ethanol until they were embedded in Paraffin. Paraffin blocks were sectioned with a 5 μm thickness. Brains were sectioned from the midline starting in a serial manner and 6 slices were collected representing a block of 600 to 700 μm thickness. The spinal cord was split in 3 – 6 parts and embedded to receive cross sections.

### Immunohistochemistry analysis in mouse tissues

The immunohistochemistry detection of IBA1 and GFAP were performed at Molecular Cytology Core Facility of Memorial Sloan Kettering Cancer Center using Discovery XT processor (Ventana Medical Systems). For IBA1: The tissue sections were blocked for 30 minutes in 10% normal goat serum.2% BSA in PBS. The primary antibody incubation (rabbit polyclonal IBA1 antibody, Wako, cat#019-19741) was used at 0.2 ug/ml. The incubation with the primary antibody was done for 5 h, followed by 60 min incubation with biotinylated goat anti-rabbit IgG (Vector labs, cat#:PK6101) in 1:200 dilution (6.5 ugr/mL). Blocker D, Streptavidin- HRP and DAB detection kit (Ventana Medical Systems) were used according to the manufacturer instructions. For GFAP: The tissue sections were blocked for 30 minutes in 10% normal goat serum.2% BSA in PBS.The primary antibody incubation (rabbit polyclonal GFAP antibody, Dako, cat#Z0334) was used at 1 ug/ml. The incubation with the primary antibody was done for 5 h, followed by 32 min incubation with biotinylated goat anti-rabbit IgG (Vector labs, cat#:PK6101),1:200 (6.5 ugr/mL). Blocker D, Streptavidin- HRP and DAB detection kit (Ventana Medical Systems) were used according to the manufacturer instructions. IBA1 and GFAP positive area was determined based on IHC staining. Sagittal brain images were segmented manually in QuPath into Cortex, Striatum, Hippocampus, Midbrain (including basal ganglia), Pons/Medulla, Spinal cord and the Cerebellum with additional separation of the white matter and the grey matter in the Cerebellum. Segmented areas and the unsegmented image were individually exported to ImageJ with a 1:4 compression. Segmented brain areas were individually analyzed in ImageJ by subtracting the background (Substract Background) with rolling ball radius of 50 pixels. Color Deconvolution 1.7 plugin was used to split the DAB channel from hematoxylin by using the H DAB vector. The DAB channel was further used and transferred into a binary image by selecting the Threshold value given by the Default or Otsu algorithm run on the unsegmented sagittal brain image for GFAP or IBA1, respectively. The DAB positive area was determined by running Analyze Particles taking all particles in the rage of 2 – Infinity Pixel into account.

### Immunofluorescence in mouse tissue

Immunofluorescence of mouse tissue was carried out on 3–4-μm thick paraffin sections, fixed with PFA with anti-pSTAT3-Tyr705 (1:100 D3A7, XP® Rabbit mAb, Cell Signaling), anti-IBA1 (1:200, AIF-1/IBA1 Antibody Goat Polyclonal antibody, Novus Biologicals) and anti-GFAP (1:200, ab4674, Chicken polyclonal, Abcam), NEUN (1:200, MAB377, Millipore). Secondary Alexa647, Alexa555 and Alexa488 (Invitrogen) were added 1:200. mages were taken with a Zeiss Laboratory.A1, BondIII (Leica-Microsystems), BZ-9000 BIOREVO microscope (Keyence) and analyzed using the BZ-II Analyzer (Keyence) or with a LSM880 Zeiss microscope with 40×/1.4 (oil), performing a tile scan and z stack on the whole tissue at a 512 × 512 or 1,024 × 1,024 pixel resolution and manually analyzed using Imaris (Bitplane) software.

### Isolation of mouse brain nuclei for sn-RNAseq

All samples were handled and processed under Air Clean PCR Workstation. Frozen brain tissue was dissociated using a Dounce homogenizer with 1X lysis buffer (Nuclei PURE Lysis Buffer, Sigma L9286). Nuclei suspension is filtered using a 35 um Cell Strainer and centrifuged at 600g for 5 min at 4 °C. Pellet was resuspended in wash buffer [1X SCC (Invitrogen, cat no AM9770) 20 mM DTT, 1% BSA and RNAse inhibitor (Ambion, cat no AM2682)] and centrifuged at 600g for 5 min at 4 °C. Nuclei were stained with DAPI at 1 mg/1 mL (Invitrogen, cat no D1306 and DAPI+ nuclei were FACS-sorted with a BD FACS Aria with a 100-μm nozzle and a sheath pressure 20 psi. Sorted nuclei were centrifuged in 5mL tubes in a swinging bucket centrifuge at 600g for 5min. RIN was determined using BioAnalyzer.

### Single-cell barcoding, library preparation and sequencing

The single-nuclei RNA-Seq of FACS-sorted nuclei suspensions was performed on Chromium instrument (10X genomics) following the user guide manual for 3′ v3.1. In brief, FACS-sorted cells were washed once with PBS containing 1% bovine serum albumin (BSA) and resuspended in PBS containing 1% BSA to a final concentration of 700–1,300 cells per μl. The viability of cells was above 80%, as confirmed with 0.2% (w/v) Trypan Blue staining (Countess II). Cells were captured in droplets. Following reverse transcription and cell barcoding in droplets, emulsions were broken and cDNA purified using Dynabeads MyOne SILANE followed by PCR amplification per manual instruction. Samples were multiplexed together on one lane of 10X Chromium (using Hash Tag Oligonucleotides - HTO) following previously published protocol ^67^. Final libraries were sequenced on Illumina NovaSeq S4 platform (R1 – 28 cycles, i7 – 8 cycles, R2 – 90 cycles). The cell-gene count matrix was constructed using the Sequence Quality Control (SEQC) package ^68^. Viable cells were identified on the basis of library size and complexity, whereas cells with >20% of transcripts derived from mitochondria were excluded from further analysis.

### Sn-RNA-seq data preprocessing

All downstream analyses were performed using AnnData v.0.7.4 and the scanpy package v.1.5.1. A matrix of 22,276 genes x 15,067 cells was used for quality control and further analysis. Cells with more than 20% of transcripts derived from mitochondria were excluded. Genes detected in fewer than ten cells or genes with low expression levels were also excluded. Mitochondrial and ribosomal genes were removed using a curated list. To remove cellular doublets, we used the DoubletDetection package with P value threshold = 1 × 10−7, voter thresh = 0.08 followed by manual inspection of the co-occurrence of contradictory markers. A matrix of 18,730 genes x 12,603 cells was retained for downstream analysis. The count matrix was normalized using the scanpy.pp.normalize total function log2-transformed using an increment of 0.1.

### Clustering analysis and cell type annotations

Principal component analysis was computed on the top 2,000 highly variable genes. Clustering was performed using PhenoGraph ^69^ and the knee point (eigenvalues smaller radius of curvature) was used to select the optimal number of principal components. For the PhenoGraph k parameter, values between k = 10 and k = 55 in steps of 5 were tested. The adjusted Rand index was used to determine the consistency of clustering across different values for k (10 to 50). The optimal k = 30 for all cell compartments was chosen from the window where the adjusted Rand index was consistently high. This approach identified N= 24 clusters. Clusters were displayed using a Uniform Manifold Approximation and Projection (UMAP). Clusters of interest were identified based on marker genes using a manually curated panel. For microglia, (CX3CR1, C1QA, PTPRC), oligodendrocytes, (MBP, MOBP), OPCs, (NEU4, PDGFRA), Astrocytes, (AQP4, GJA1), Neurons, (SYT1, SNAP25, GRN1), Vascular/Fibroblasts, (DCN, VTN), T cells, (CD3D, CD52). Each cell type of interest was re-clustered as described above and analyzed separately.

### Annotation of neuronal clusters

The identities of neuron clusters from brainstem snRNA-seq were annotated using Mousebrain.org (http://mousebrain.org/adolescent/genesearch.html). The top 10 unique genes from each cluster were searched as a group within the adolescent brain atlas and best match was identified based on magnitude of expression, number of genes expressed, and anatomical annotation to the hindbrain. Neuron identities were further confirmed by examining anatomical localization of top gene expression via Spatial RNAseq data of adult brains from Mousebrain.org (http://mousebrain.org/adult/spatial.html) and in situ hybridization data from Allen Brain Atlas (https://mouse.brain-map.org/search/index).

### Differential gene expression

Differential expression analysis between the PhenoGraph-generated clusters was performed using MAST ^70^ with default parameters. Differentially expressed genes were considered statistically significant if they had an FDR <= 0.05 and the absolute value of the log2(fold change) was greater or equal to 0.5.

### Gene set enrichment

Lists of qualified differentially expressed genes between clusters within different cell types were used for pathway analysis using the enrichr module from the gseapy package. Gene sets and pathways included, GO_Biological_Process_2018, MSigDB_Hallmark_2020, KEGG_2019_Mouse and Reactome_2016. Results with FDR<0.05 were reported as significantly enriched pathways. Gene signatures of reactive astrocytes were obtained from ^71^ ^72^. Scoring of gene signatures across clusters was done by applying the sc.tl.score_genes function from scapy on the clusters of interest. Briefly, the score is the average log-transformed expression of a set of genes subtracted with the average log-transformed expression of a reference set of genes.

### Mouse drug treatments

For BRAF and/or CSF1R inhibition, mice were placed on an ad libitum PLX4720 (BRAF inhibitor), PLX5622 (CSF1R inhibitor), or combined diet at three months of age. PLX4720 chow (417 mg/kg) and PLX5622 chow (1,200 mg/kg) were provided by Plexxikon Inc)^37,39^. Braf WT and Braf VE male and female littermates were assigned randomly into the control or treated group. Scoring of mice was performed blinded weekly by assessing hindlimb reflexes and other behavioral phenotypes such as axial rolling, as described previously ^25^. The investigators were not blinded to allocation during experiments and outcome assessment.

### Statistical analysis

Statistical analyses were performed with GraphPad Prism (v8) and R (3.6.3). Tests used are detailed in corresponding figure legends.

## Supplemental information titles and legends

### Supplemental Tables

Table S1: List of control and histiocytosis patients analyzed.

Table S2: List of genes in the HemePACT panel

Table S3: List of somatic mutations identified by HemePACT in brain nuclei from patients and controls, with ddPCR validation.

Table S4: RNA-seq analysis of human whole brain samples. Differentially Regulated Genes and Pathway Analysis.

Table S5: Bulk RNA-seq analysis of sorted microglia from mouse cortex and brainstem at 2 months and end stage of the disease. Differentially Regulated Genes and pathway analysis.

Table S6: Conserved gene signature between human brain cells and mouse microglia. Common genes and corresponding pathway analysis.

Table S7: sn-RNAseq. Differential Expression Analysis between Microglia Cluster 4 and Cluster 0,1,3

Table S8: sn-RNAseq. Differential Expression Analysis and Pathway Analysis between Astrocytes Cluster 3 and Cluster 2 and 1.

Table S9: sn-RNAseq: Differential Expression Analysis and Pathway Analysis between Oligodendrocytes Cluster 2, 4 and 5 and Cluster 0 and 1.

### Supplementary Figures

**Figure S1:**
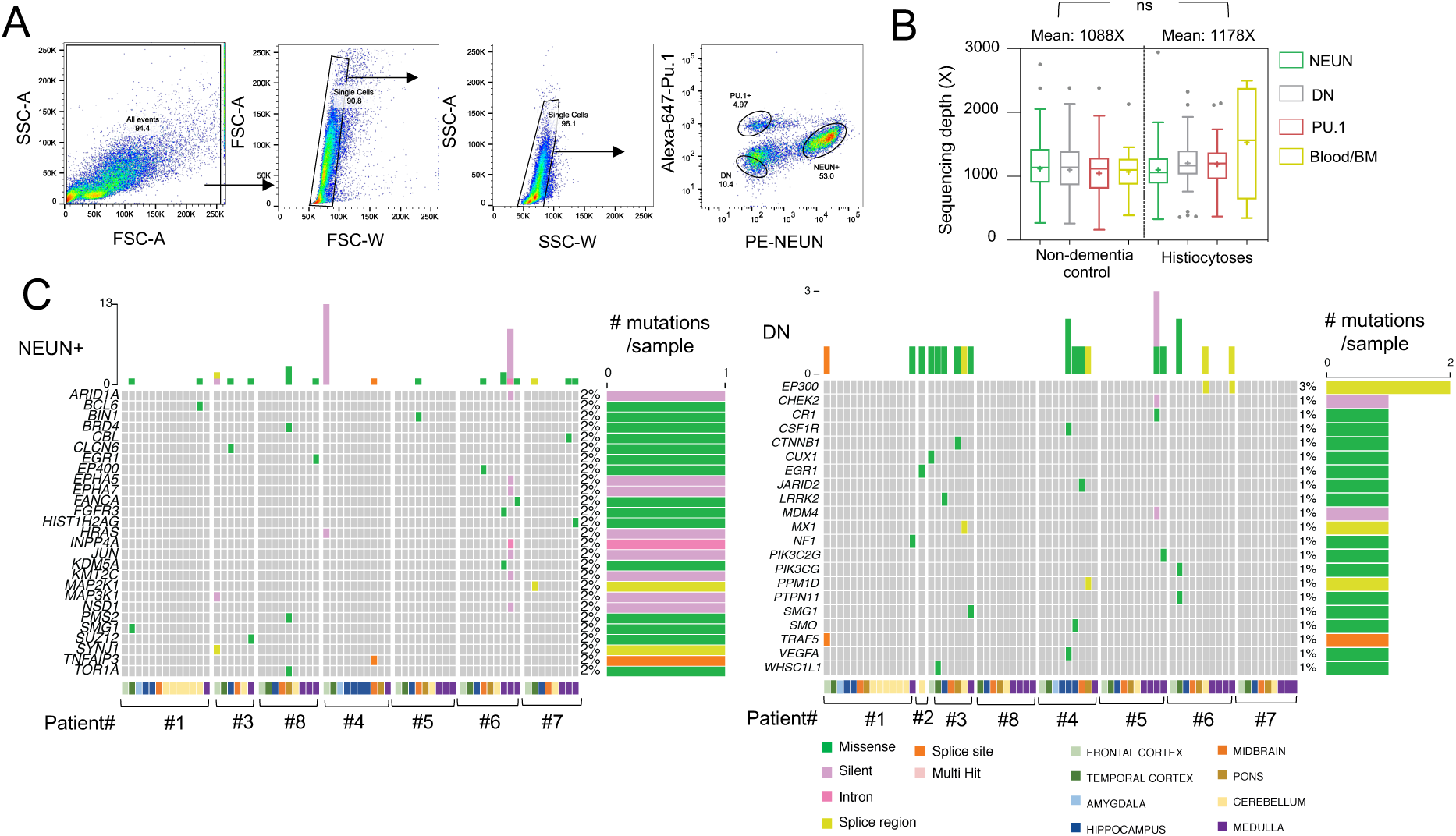
Detection of mutations in FACS-sorted brain cell types and blood PBMCs from histiocytosis patients and controls. **(A)** Flow cytometry plots illustrate the gating strategy for nuclei sorting used to separate PU.1+, NeuN+ and DN nuclei. Sorting purity was >90%. **(B)** Boxplot represents HemePACT sequencing depth (median: line, mean: cross, 25-75^th^ quartiles (boxes) and minimum/maximum (whiskers) for PU.1+ nuclei, PU.1+, NeuN+, and negative nuclei (DN, NeuN-. PU.1-) in brain samples from ‘healthy’ control brains (151 brain samples from 35 individuals, see Vicario et al 2023b) and histiocytosis (71 brain samples from 8 patients). Statistics: p value was calculated with Anova, ns: not significant. **(C)** Oncoplots represent all genes carrying mutations detected by targeted sequencing (HemePACT, Table S3) in NeuN+ sorted samples (n=71) and DN sorted samples (n=71) from histiocytosis patients (n=8). Left, number of mutations per sample and right % of samples carrying mutations.

**Figure S2:**
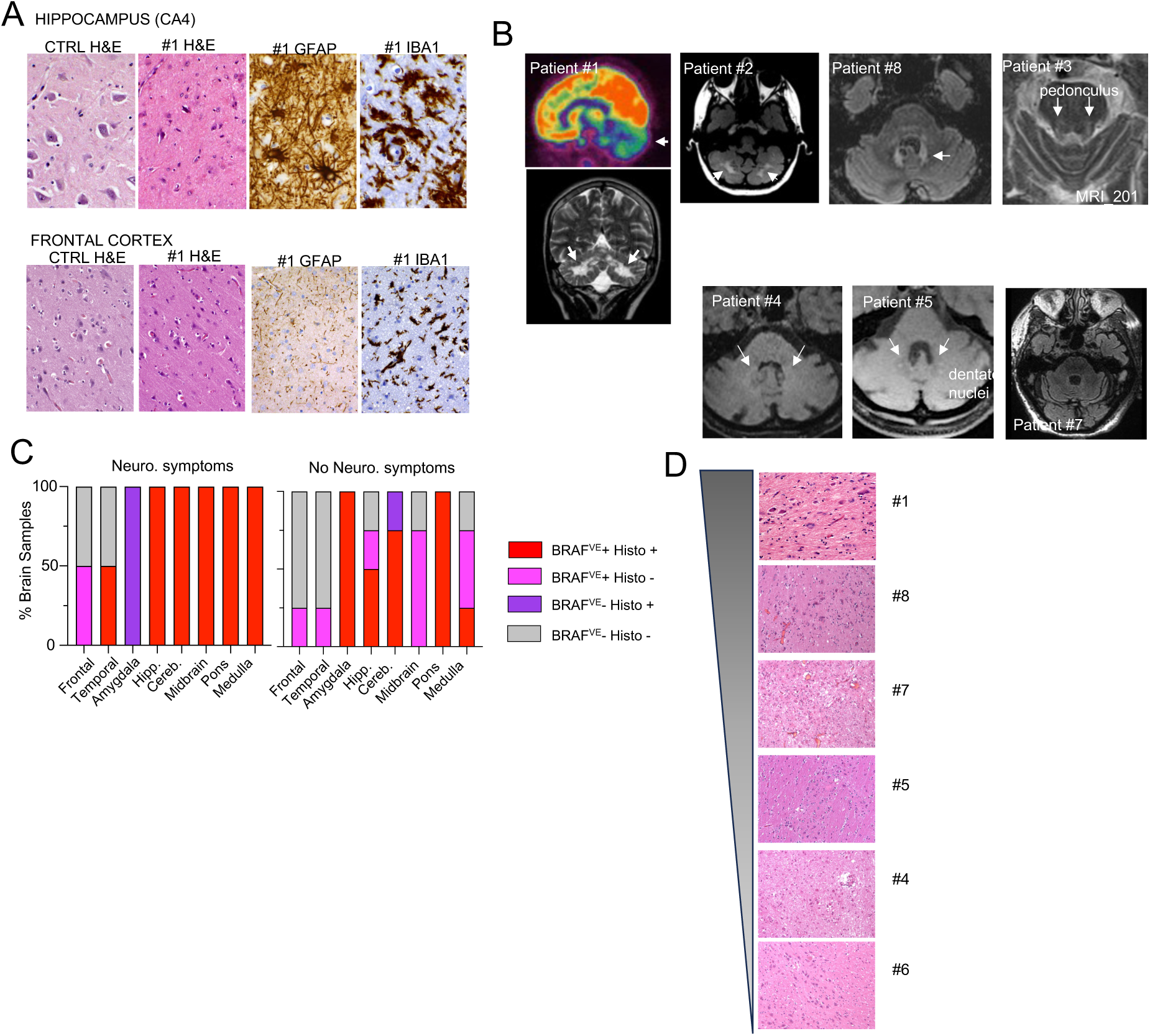
Histological and molecular analysis of histiocytosis patients. **(A)** H&E, IBA1 (microglia marker) and GFAP (astrocyte marker) staining of hippocampus and frontal cortex from patient #1 and an age-matched control individual for comparison. **(B)** For patient #1, PET scan (top) at age 9 shows hypometabolism in the cerebellum (arrow) and the thalamic region MRI at age 18 (bottom) shows hyperintensity signals in pons, dentate nuclei, and the cerebellum (arrows). MRI images show hyperintensity signals in the cerebellum dentate nuclei and cerebellar pedunculus of patient 2,3,8,4 and 5 (arrows). Patient 7 MRI did not reveal detectable abnormalities. No MRI was available for patient 6. **(C)** Bar graphs represent the proportion of tested brain samples positive for BRAFc.1799T>A (V600E) by droplet-digital PCR (ddPCR) and/or histological signs of neurodegeneration among patients with (left) or without (right) neuro-histiocytosis. **(D)** Representative H&E from pons and cerebellum from patients ranked by neuronal damage by two pathologists, from higher to lower.

**Figure S3:**
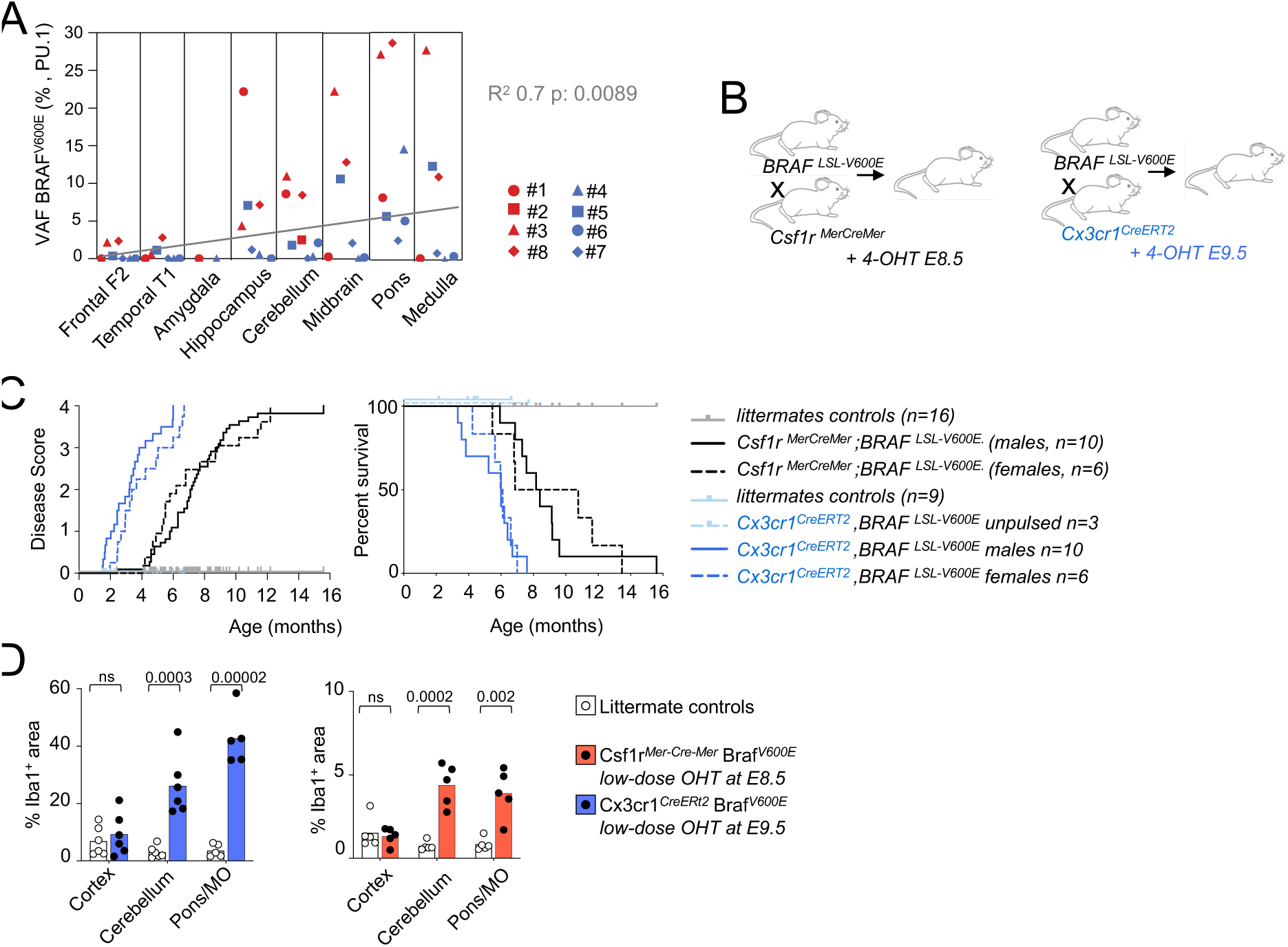
Mouse models of histiocytosis. **(A)** Variant allelic frequency (VAF, %, by ddPCR) of BRAFc.1799T>A (V600E) in PU.1+ nuclei from histiocytosis patients across brain regions (n=8, patients with neuro-histiocytosis are color-coded in red, patients without a diagnosis of neuro-histiocytosis are color-coded in blue). The fitted line, R-squared and corresponding p value were calculated by simple linear regression by assigning numbers from 1-8 to each brain region from along a rostro caudal axis. **(B)** Mouse models of neuro-histiocytosis. *Csf1r^MerCreMer^* and *Cx3cr1^CreERt2^* pregnant females crossed with *Braf^LSL-V600E^* males receive low-dose 4-Hydroxytamoxifen (a single injection of 37.5 mg 4-OHT per kg, ip, and 18.75 mg per kg of body weight of progesterone) at E8.5 and E9.5 respectively. **(C)** Plots represent disease score progression and survival over time of mice in B. **(D)** Quantification of % of IBA1+ area from *Csf1r^MerCreMer^* Braf*^LSL-V600E^* and *Cx3cr1^CreERt2^* Braf*^LSL-V600E^* mice and littermate controls. Statistics: p values were calculated with unpaired t-test with Welch’s correction.

**Figure S4:**
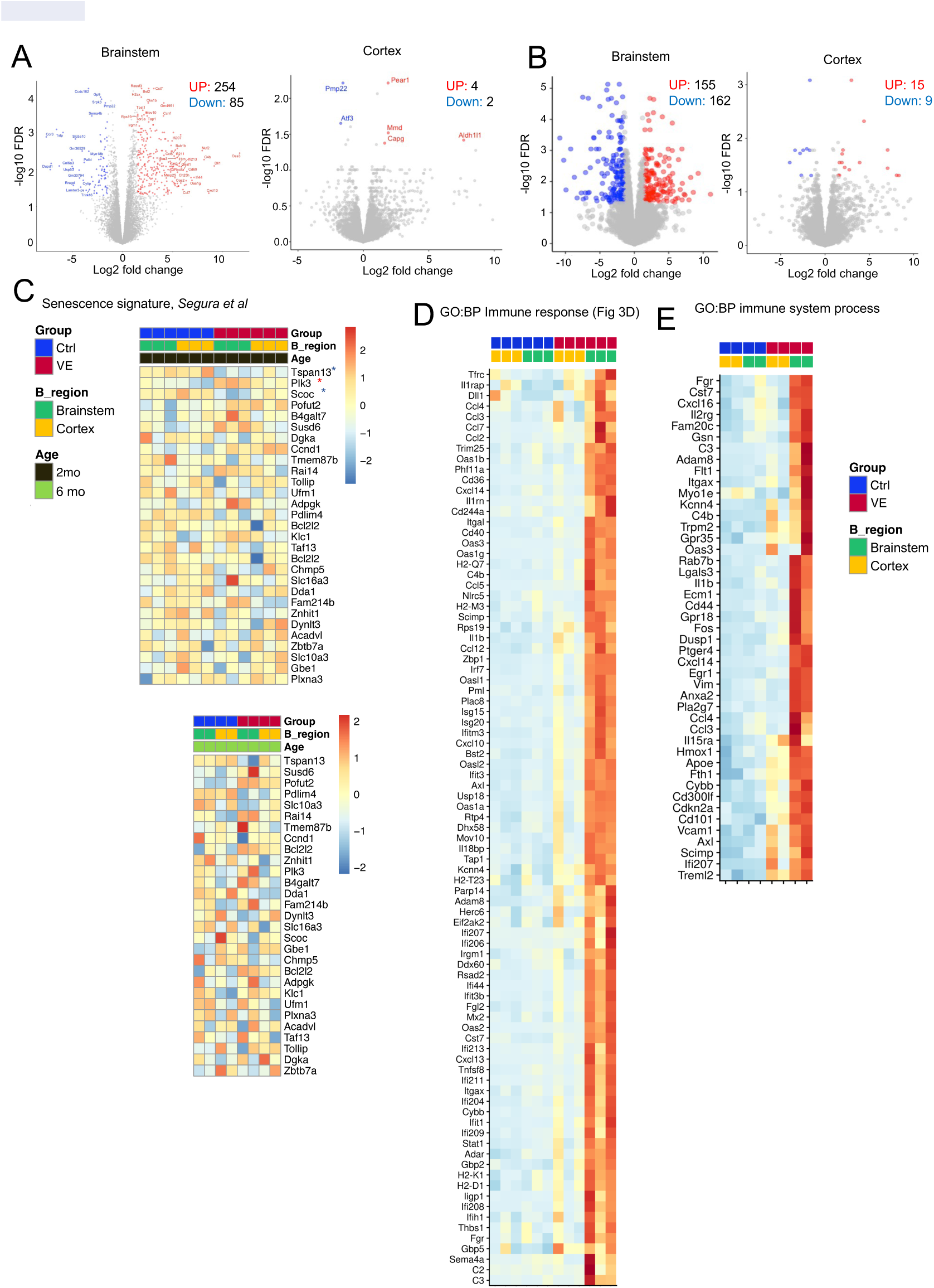
Bulk transcriptional analysis of mouse microglia. **(A)** Volcano plots of differentially expressed genes from FACS-sorted microglia from 2-month-old *Cx3cr1^CreERt2^* Braf*^LSL-V600E^* mice versus microglia from control mice in brainstem (left) and in cortex (right). X-axis represents the significance (−Log10(p-value)) and Y-axis represents the fold-change (Log2(fold-change)). Upregulated genes (log2FC >=1.5 & FDR <=0.05) are represented with red dots, downregulated genes (log2FC <= −1.5 & FDR <=0.05) are represented with blue dots. **(B)** Volcano plots of differentially expressed genes from FACS-sorted microglia from ∼6 month old (end stage) *Cx3cr1^CreERt2^* Braf*^LSL-V600E^* mice versus microglia from control mice in brainstem (left) and in cortex (right). X-axis represents the significance (−Log10(p-value)) and Y-axis represents the fold-change (Log2(fold-change)). Upregulated genes (log2FC >=1.5 & FDR <=0.05) are represented with red dots, downregulated genes (log2FC <= −1.5 & FDR <=0.05) are represented with blue dots. **(C)** Hierarchical clustering of genes from reported senescence signature from Segura et al. in 2-month-old *Cx3cr1^CreERt2^* Braf*^LSL-V600E^* mice (top) and 6 month old (end stage)(bottom) and littermates controls. Note: genes with red star indicate significantly regulated. **(D)** Hierarchical clustering of DEG from ‘GO:BP Immune response’ (GO:0006955, left) and GO:BP ‘immune system process (GO:0002376, right) from analysis in Figure 3D.

**Figure S5.**
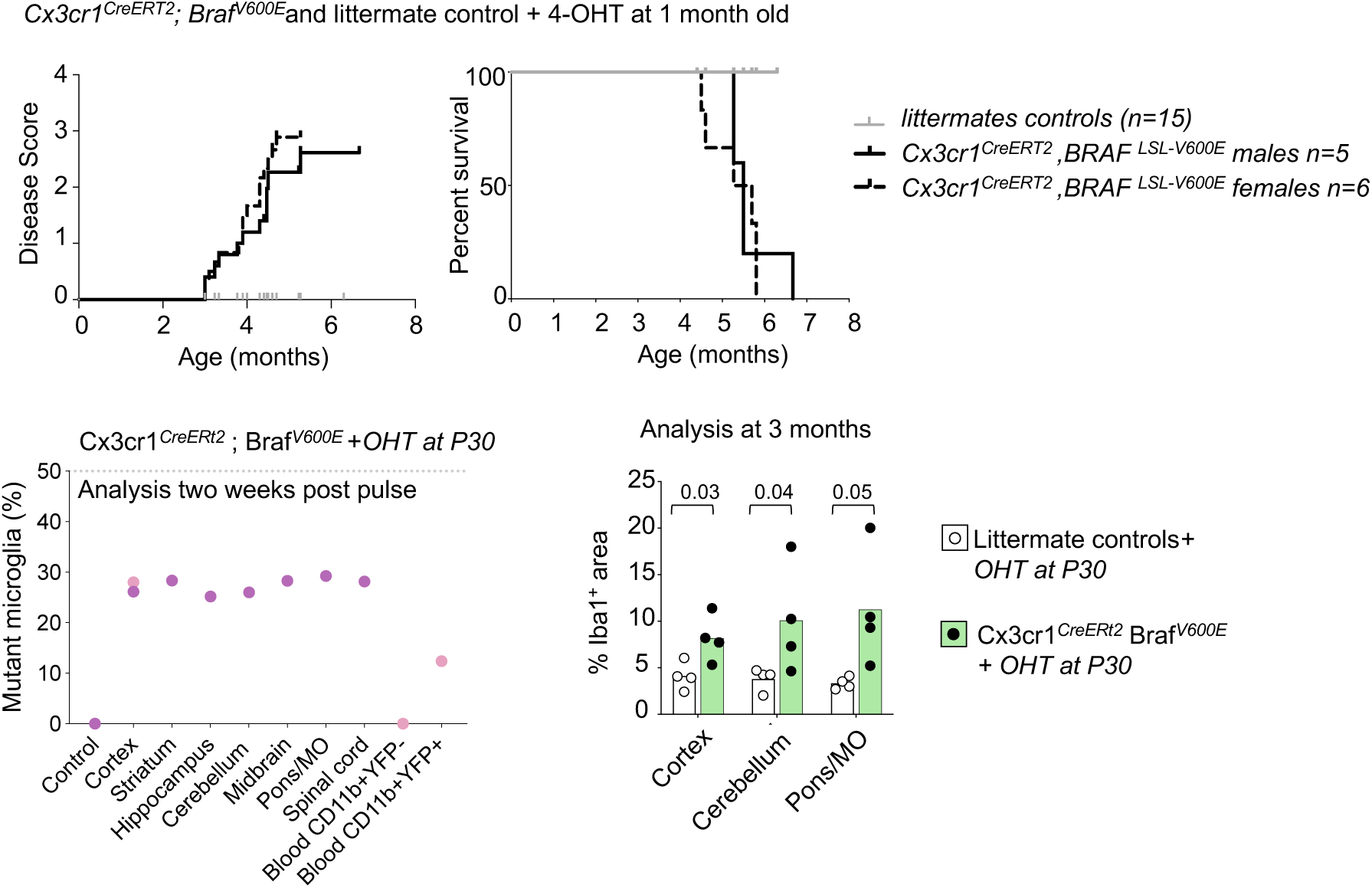
Mouse models of histiocytosis. One month old *Cx3cr1^CreERt2^*; *Braf^LSL-V600E^* mice were pulsed with high-dose 4-OHT (100 mg/kg Tamoxifen per kg/day for 5 days, intra peritoneally). Top, plots represent disease score progression and survival over time. Bottom left, allelic frequency of recombined mutant *Braf* allele by digital PCR in microglia sort-purified from different brain regions from mice in B, two weeks after 4-OHT injection. Right, quantification of % of IBA1+ area. Statistics: p values were calculated with unpaired t-test with Welch’s correction.

**Figure S6:**
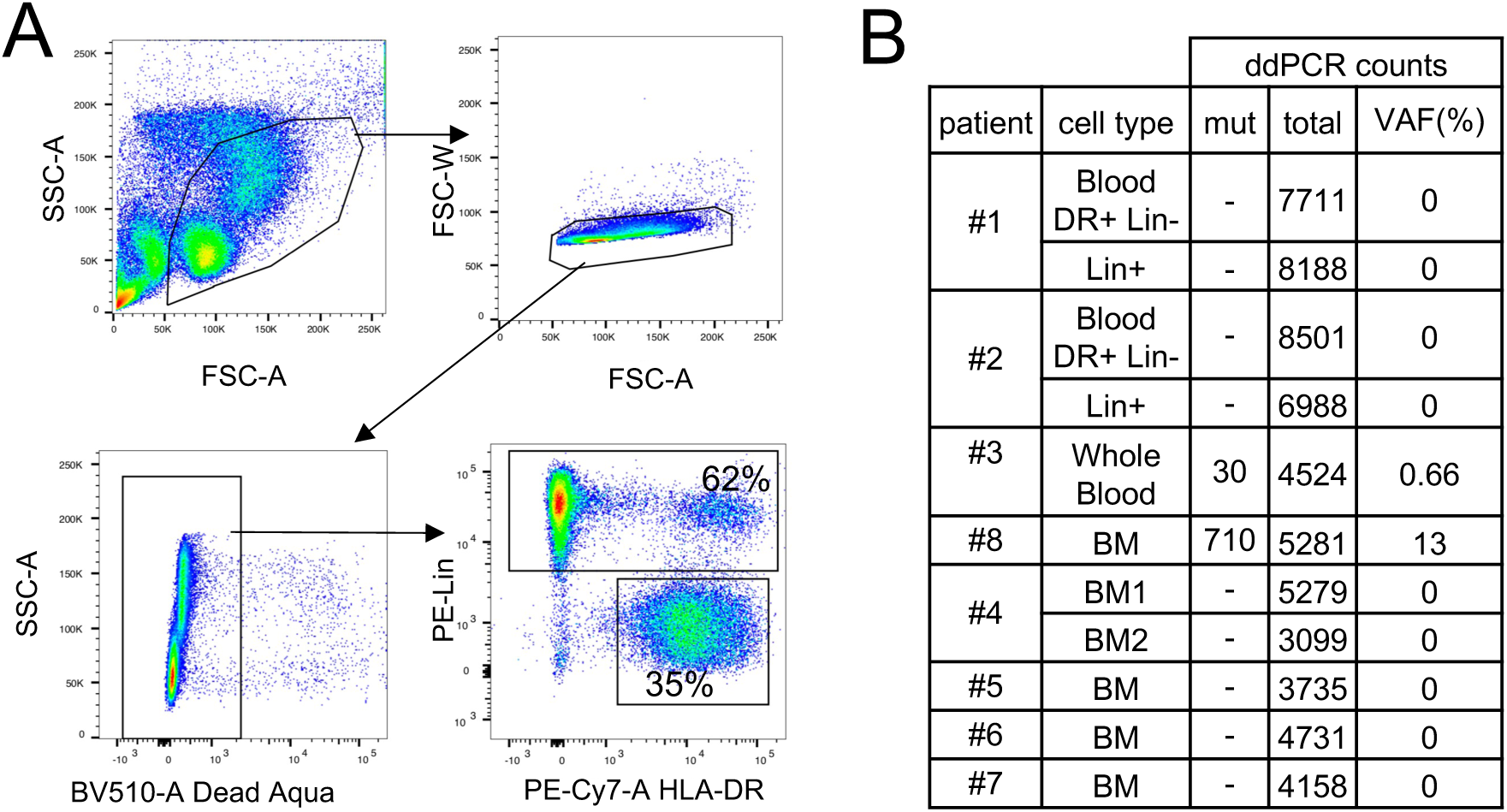
Analysis of Blood and Bone Marrow from histiocytosis patients. **(A)** Flow cytometry plots illustrate the gating strategy used to separate monocyte/DCs (HLA-DR+, CD3-/CD15-/CD19-/CD56-/NKp46-(Lin) and Lin+ cells for patients #1 and 2. (B) BRAFV600E detection by ddPCR from blood or bone marrow from Histiocytosis patients. Mutant count>5 are considered positive, Mut: mutant counts. VAF: Variant Allelic Frequency (%).

**Figure S7:**
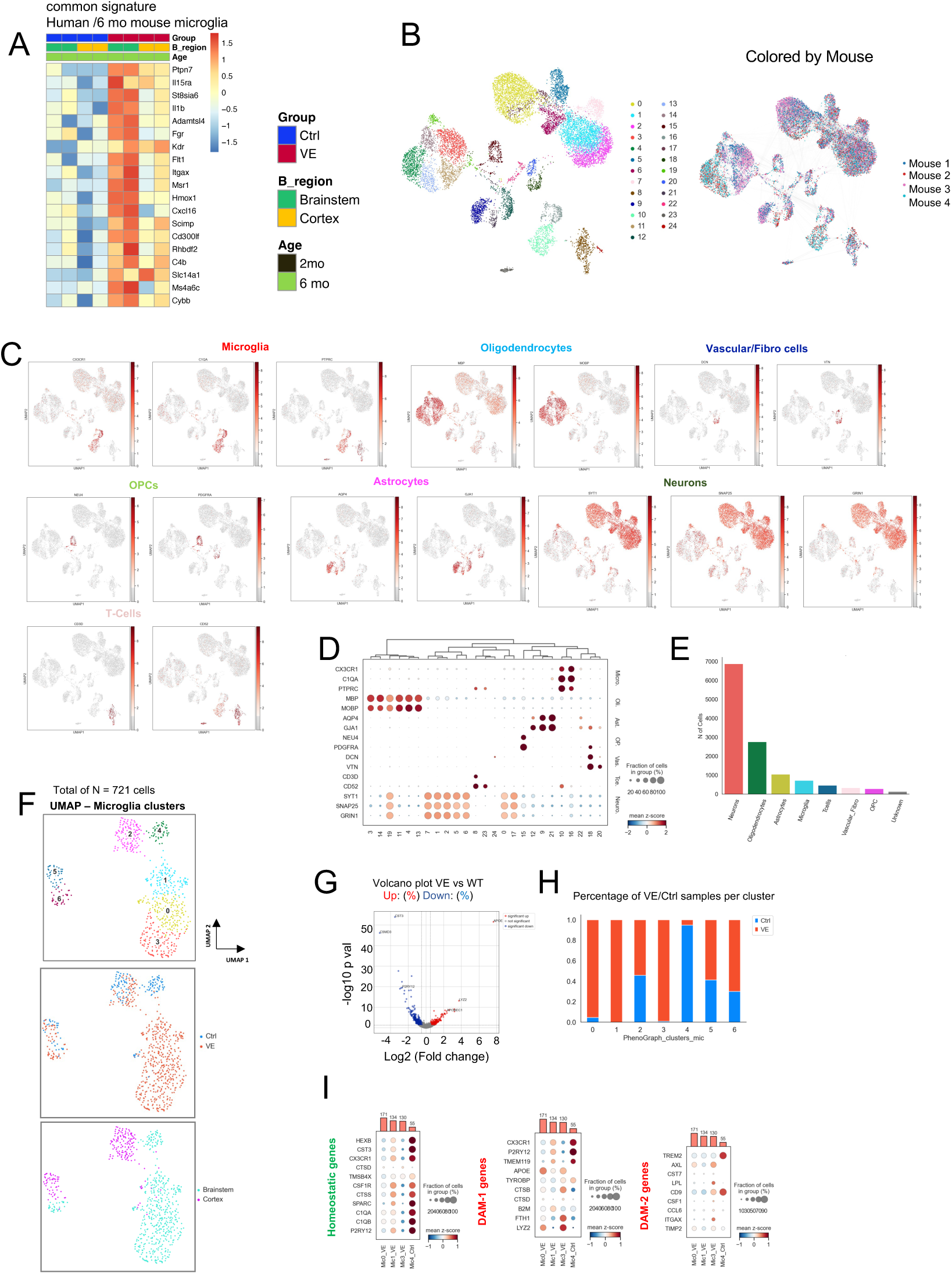
Transcriptomic analysis from human and mouse brain samples. **(A)** Hierarchical clustering of common genes in end stage mouse microglia from *Cx3cr1^CreERt2^* Braf*^LSL-V600E^* in Figure 3 and human whole brain samples in Figure 2. Expression values are *Z* score transformed. Samples were clustered using average linkage and cluster similarity was determined using the Euclidean distance **(B) sn-RNAseq**. Left, UMAP of single nuclei RNA profiles from Cx3cr1^CreER^ *Braf^LSL-V600E^* mice pulsed with OH-TAM at E9.5 and analyzed at 6 month of age (end stage, VE) (n=2) and littermate controls (Ctrl, n=2) on n=12,603 nuclei depicting n=24 clusters. Right, UMAP depicting separation of nuclei by mouse. **(C)** Expression of marker genes across clusters. UMAP, colored by expression levels of marker genes: Microglia (CX3CR1, C1QA, PTPRC), Oligodendrocytes (MBP, MOBP), Vascular/Fibroblasts (DCN, VTN), OPCs (NEU4, PDGFRA), Astrocytes (AQP4, GJA1), Neurons (SYT1, SNAP25, GRIN1), Tcells (CD3D, CD52). **(D)** Dot plot showing the expression level (color scale) and the percent of cells expressing (dot size) marker genes across all clusters (rows). **(E)** Barplot showing the numbers of cells classified as Microglia, Oligodendrocytes, Vascular/Fibroblasts, OPCs, Astrocytes, Neurons and T cells out of the total nuclei population (n =12,603). **(F)** UMAP of sn-RNAseq profiles of identified microglia (n = 721) from *Cx3cr1^CreER^ Braf^LSL-V600E^* and control mice. Clusters are colored by cluster assignment, condition, and brain region, respectively. **(G)** Volcano plot of differentially expressed genes between microglia clusters (Mic0, Mic1, Mic3 vs Mic4). X-axis represents the significance (−Log10(p-value)) and Y-axis represents the fold-change (Log2(fold-change)). Upregulated genes (log2FC >=0.5 & p-value <=0.05) are represented with red dots, downregulated genes (log2FC <= −0.5 & p-value <=0.05) are represented with blue dots. **(H)** Stacked bar chart of percentage contribution of each condition across clusters of the microglia population. **(I)** Dot plot showing the expression level (color scale) and the percent of cells expressing (dot size) (left) homeostatic microglia markers, (center) DAM-1 markers and (right) DAM-1 markers across all clusters (rows). Barplot at the top represents number of cells per cluster.

**Figure S8:**
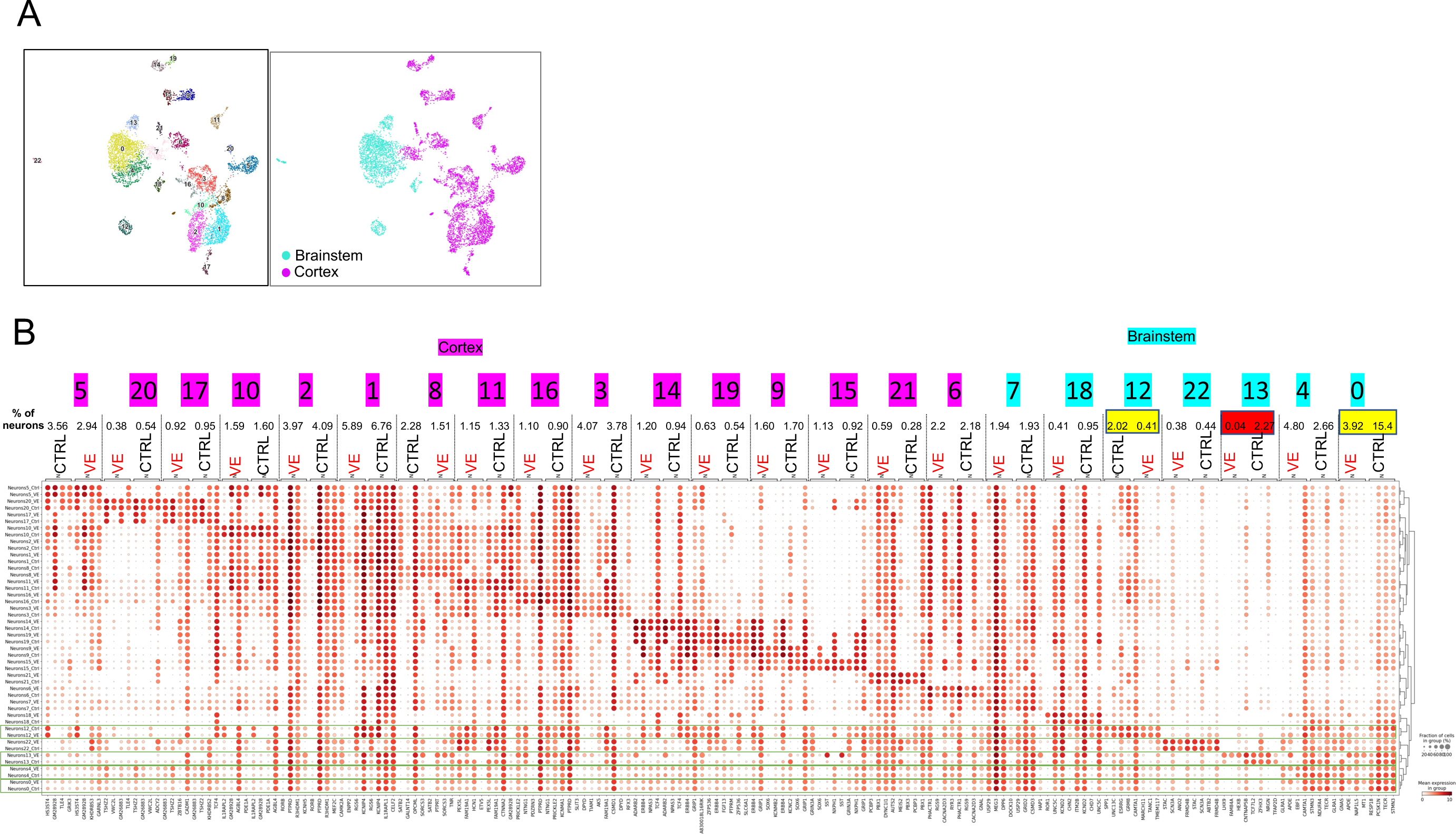
Sn-RNAseq from mouse brain samples: neuronal clusters. **(A)** UMAP representation of neurons (n=6,871). Clusters are colored by cluster assignment. **(B)** Dot plot showing the expression level (color scale) and the percent of cells expressing (dot size) of top expressed genes in each neuronal control cluster vs the rest of control clusters in *Cx3cr1^CreER^ Braf^LSL-V600E^*and control mice.

**Figure S9:**
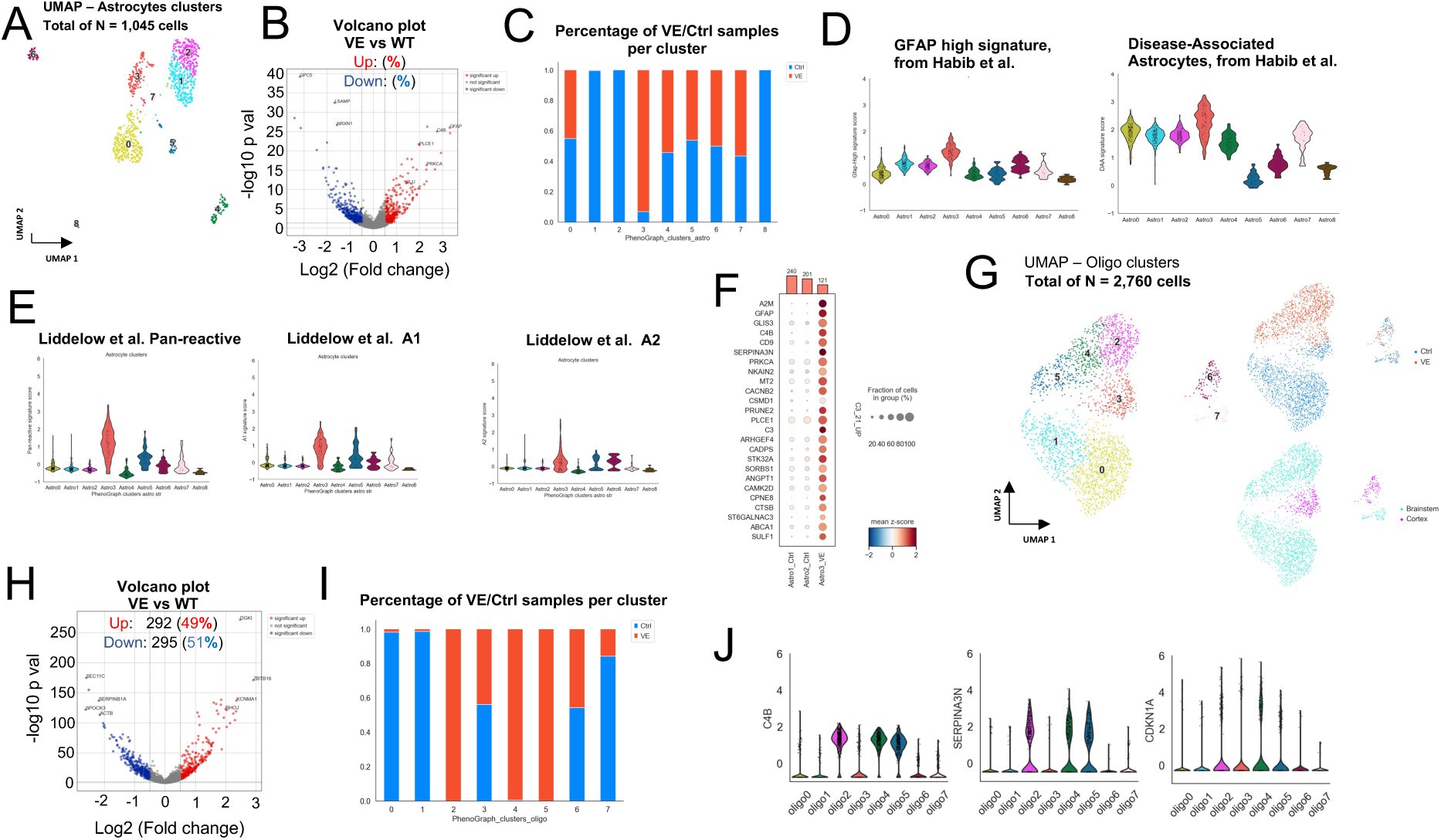
sn-RNAseq from mouse brain samples: differentially expressed genes in astrocytes and oligodendrocytes. **(A)** UMAP representation of identified astrocytes (n=1,045). Clusters are colored by cluster assignment. **(B)** Volcano plot of differentially expressed genes between astrocyte clusters (Astro3 (VE) vs Astro2, Astro1 (WT)). X-axis represents the significance (−Log10(p-value)) and Y-axis represents the fold-change (Log2(fold-change)). Upregulated genes (log2FC >=0.5 & p-value <=0.05) are represented with red dots, downregulated genes (log2FC <= −0.5 & p-value <=0.05) are represented with blue dots. **(C)** Stacked bar chart of percentage contribution of each condition across clusters of the astrocyte population. **(D, E)** Signatures of reactive astrocytes. Violin plots show the distribution of gene expression scores across all clusters of identified astrocytes in previously defined signatures. **(F)** Dot plot showing the expression level (color scale) and the percent of cells expressing (dot size) the top 20 most significantly upregulated genes (log2FC >= 0.5 & FDR <= 0.05) between astrocytes A3 (VE) vs A1, A2 (control). Bar plot at the top represents number of cells per cluster. **(G)** UMAP representation of identified oligodendrocytes (n=2,760). Clusters are colored by cluster assignment, condition, or brain region. **(H)** Volcano plot of differentially expressed genes between oligodendrocytes clusters (O2, O4, O5 (VE) vs O0, O1 (WT)). X-axis represents the significance (−Log10(p-value)) and Y-axis represents the fold-change (Log2(fold-change)). Upregulated genes (log2FC >=0.5 & p-value <=0.05) are represented with red dots, downregulated genes (log2FC <= −0.5 & p-value <=0.05) are represented with blue dots. **(I)** Stacked bar chart of percentage contribution of each condition across clusters of the oligodendrocyte population. **(J)** Expression of DA1 Oligodendrocyte markers C4B, SERPINA3N and CDKN1A across oligodendrocyte clusters.

## References

1. Heritier S, Emile JF, Barkaoui MA, et al. BRAF Mutation Correlates With High-Risk Langerhans Cell Histiocytosis and Increased Resistance to First-Line Therapy. J Clin Oncol 2016;34(25):3023–30. DOI: 10.1200/JCO.2015.65.9508.

2. Haroche J, Charlotte F, Arnaud L, et al. High prevalence of BRAF V600E mutations in Erdheim-Chester disease but not in other non-Langerhans cell histiocytoses. Blood 2012;120(13):2700–3. DOI: 10.1182/blood-2012-05-430140.

3. Boyd LC, O’Brien KJ, Ozkaya N, et al. Neurological manifestations of Erdheim-Chester Disease. Ann Clin Transl Neurol 2020;7(4):497–506. DOI: 10.1002/acn3.51014.

4. Rigaud C, Barkaoui MA, Thomas C, et al. Langerhans cell histiocytosis: therapeutic strategy and outcome in a 30-year nationwide cohort of 1478 patients under 18 years of age. Br J Haematol 2016;174(6):887–98. DOI: 10.1111/bjh.14140.

5. Emile JF, Abla O, Fraitag S, et al. Revised classification of histiocytoses and neoplasms of the macrophage-dendritic cell lineages. Blood 2016;127(22):2672–81. DOI: 10.1182/blood-2016-01-690636.

6. Badalian-Very G, Vergilio JA, Degar BA, et al. Recurrent BRAF mutations in Langerhans cell histiocytosis. Blood 2010;116(11):1919–23. DOI: 10.1182/blood-2010-04-279083.

7. Satoh T, Smith A, Sarde A, et al. B-RAF mutant alleles associated with Langerhans cell histiocytosis, a granulomatous pediatric disease. PLoS One 2012;7(4):e33891. DOI: 10.1371/journal.pone.0033891.

8. Durham BH, Lopez Rodrigo E, Picarsic J, et al. Activating mutations in CSF1R and additional receptor tyrosine kinases in histiocytic neoplasms. Nat Med 2019;25(12):1839–1842. DOI: 10.1038/s41591-019-0653-6.

9. Cohen Aubart F, Idbaih A, Galanaud D, et al. Central nervous system involvement in Erdheim-Chester disease: An observational cohort study. Neurology 2020;95(20):e2746–e2754. (In eng). DOI: 10.1212/wnl.0000000000010748.

10. Heritier S, Barkaoui MA, Miron J, et al. Incidence and risk factors for clinical neurodegenerative Langerhans cell histiocytosis: a longitudinal cohort study. Br J Haematol 2018;183(4):608–617. DOI: 10.1111/bjh.15577.

11. Bhatia A, Hatzoglou V, Ulaner G, et al. Neurologic and oncologic features of Erdheim-Chester disease: a 30-patient series. Neuro Oncol 2020. DOI: 10.1093/neuonc/noaa008.

12. Wnorowski M, Prosch H, Prayer D, Janssen G, Gadner H, Grois N. Pattern and course of neurodegeneration in Langerhans cell histiocytosis. J Pediatr 2008;153(1):127–32. DOI: 10.1016/j.jpeds.2007.12.042.

13. Barthez MA, Araujo E, Donadieu J. Langerhans cell histiocytosis and the central nervous system in childhood: evolution and prognostic factors. Results of a collaborative study. J Child Neurol 2000;15(3):150–6. DOI: 10.1177/088307380001500302.

14. Prosch H, Grois N, Wnorowski M, Steiner M, Prayer D. Long-term MR imaging course of neurodegenerative Langerhans cell histiocytosis. AJNR Am J Neuroradiol 2007;28(6):1022–8. DOI: 10.3174/ajnr.A0509.

15. Laurencikas E, Gavhed D, Stalemark H, van’t Hooft I, Prayer D, Grois N, Henter JI. Incidence and pattern of radiological central nervous system Langerhans cell histiocytosis in children: a population based study. Pediatr Blood Cancer 2011;56(2):250–7. DOI: 10.1002/pbc.22791.

16. Papo M, Emile JF, Maciel TT, et al. Erdheim-Chester Disease: a Concise Review. Curr Rheumatol Rep 2019;21(12):66. DOI: 10.1007/s11926-019-0865-2.

17. Grois N, Prayer D, Prosch H, Lassmann H, Group CLC-o. Neuropathology of CNS disease in Langerhans cell histiocytosis. Brain 2005;128(Pt 4):829–38. DOI: 10.1093/brain/awh403.

18. McClain KL, Picarsic J, Chakraborty R, et al. CNS Langerhans cell histiocytosis: Common hematopoietic origin for LCH-associated neurodegeneration and mass lesions. Cancer 2018;124(12):2607–2620. DOI: 10.1002/cncr.31348.

19. Collin M, Bigley V, McClain KL, Allen CE. Cell(s) of Origin of Langerhans Cell Histiocytosis. Hematol Oncol Clin North Am 2015;29(5):825–38. (In eng). DOI: 10.1016/j.hoc.2015.06.003.

20. Cohen Aubart F, Roos-Weil D, Armand M, et al. High frequency of clonal hematopoiesis in Erdheim-Chester disease. Blood 2021;137(4):485–492. (In eng). DOI: 10.1182/blood.2020005101.

21. Wilk CM, Cathomas F, Torok O, et al. Circulating senescent myeloid cells infiltrate the brain and cause neurodegeneration in histiocytic disorders. Immunity 2023;56(12):2790–2802 e6. DOI: 10.1016/j.immuni.2023.11.011.

22. Schulz C, Gomez Perdiguero E, Chorro L, et al. A lineage of myeloid cells independent of Myb and hematopoietic stem cells. Science 2012;336(6077):86-90. DOI: 10.1126/science.1219179.

23. Gomez Perdiguero E, Klapproth K, Schulz C, et al. Tissue-resident macrophages originate from yolk-sac-derived erythro-myeloid progenitors. Nature 2015;518(7540):547–51. DOI: 10.1038/nature13989.

24. Mass E, Ballesteros I, Farlik M, et al. Specification of tissue-resident macrophages during organogenesis. Science 2016;353(6304). DOI: 10.1126/science.aaf4238.

25. Mass E, Jacome-Galarza CE, Blank T, et al. A somatic mutation in erythro-myeloid progenitors causes neurodegenerative disease. Nature 2017;549(7672):389–393. DOI: 10.1038/nature23672.

26. Evrony GD, Cai X, Lee E, et al. Single-neuron sequencing analysis of L1 retrotransposition and somatic mutation in the human brain. Cell 2012;151(3):483–96. DOI: 10.1016/j.cell.2012.09.035.

27. Cheng DT, Mitchell TN, Zehir A, et al. Memorial Sloan Kettering-Integrated Mutation Profiling of Actionable Cancer Targets (MSK-IMPACT): A Hybridization Capture-Based Next-Generation Sequencing Clinical Assay for Solid Tumor Molecular Oncology. J Mol Diagn 2015;17(3):251–64. DOI: 10.1016/j.jmoldx.2014.12.006.

28. Tay TL, Mai D, Dautzenberg J, et al. A new fate mapping system reveals context-dependent random or clonal expansion of microglia. Nature Neuroscience 2017;20(6):793–803. DOI: 10.1038/nn.4547.

29. Vicario R, Fragkogianni S, Weber L, et al. A microglia clonal inflammatory disorder in Alzheimer’s Disease. bioRxiv 2024. DOI: 10.1101/2024.01.25.577216.

30. Askew K, Li K, Olmos-Alonso A, et al. Coupled Proliferation and Apoptosis Maintain the Rapid Turnover of Microglia in the Adult Brain. Cell Rep 2017;18(2):391–405. DOI: 10.1016/j.celrep.2016.12.041.

31. Reu P, Khosravi A, Bernard S, et al. The Lifespan and Turnover of Microglia in the Human Brain. Cell Rep 2017;20(4):779–784. DOI: 10.1016/j.celrep.2017.07.004.

32. Ginhoux F, Greter M, Leboeuf M, et al. Fate mapping analysis reveals that adult microglia derive from primitive macrophages. Science 2010;330(6005):841–5. DOI: 10.1126/science.1194637.

33. Genovese G, Kahler AK, Handsaker RE, et al. Clonal hematopoiesis and blood-cancer risk inferred from blood DNA sequence. N Engl J Med 2014;371(26):2477–87. DOI: 10.1056/NEJMoa1409405.

34. Jaiswal S, Fontanillas P, Flannick J, et al. Age-related clonal hematopoiesis associated with adverse outcomes. N Engl J Med 2014;371(26):2488–98. DOI: 10.1056/NEJMoa1408617.

35. Ben Haim L, Ceyzériat K, Carrillo-de Sauvage MA, et al. The JAK/STAT3 pathway is a common inducer of astrocyte reactivity in Alzheimer’s and Huntington’s diseases. J Neurosci 2015;35(6):2817–29. (In eng). DOI: 10.1523/jneurosci.3516-14.2015.

36. Lee S-H, Rezzonico MG, Friedman BA, et al. TREM2-independent oligodendrocyte, astrocyte, and T cell responses to tau and amyloid pathology in mouse models of Alzheimer disease. Cell Reports 2021;37(13):110158. DOI: 10.1016/j.celrep.2021.110158.

37. Spangenberg E, Severson PL, Hohsfield LA, et al. Sustained microglial depletion with CSF1R inhibitor impairs parenchymal plaque development in an Alzheimer’s disease model. Nat Commun 2019;10(1):3758. DOI: 10.1038/s41467-019-11674-z.

38. Diamond EL, Francis JH, Lacouture ME, et al. CSF1R inhibition for histiocytic neoplasm with CBL mutations refractory to MEK1/2 inhibition. Leukemia 2023;37(8):1737–1740. DOI: 10.1038/s41375-023-01947-4.

39. Tsai J, Lee JT, Wang W, et al. Discovery of a selective inhibitor of oncogenic B-Raf kinase with potent antimelanoma activity. Proc Natl Acad Sci U S A 2008;105(8):3041–6. DOI: 10.1073/pnas.0711741105.

40. Bouzid H, Belk JA, Jan M, et al. Clonal hematopoiesis is associated with protection from Alzheimer’s disease. Nature Medicine 2023. DOI: 10.1038/s41591-023-02397-2.

41. Menassa DA, Muntslag TAO, Martin-Estebané M, et al. The spatiotemporal dynamics of microglia across the human lifespan. Developmental Cell 2022;57(17):2127–2139.e6. DOI: 10.1016/j.devcel.2022.07.015.

42. Friedman BA, Srinivasan K, Ayalon G, et al. Diverse brain myeloid expression profiles reveal distinct microglial activation states and aspects of Alzheimer’s disease not evident in mouse models. Cell reports 2018;22(3):832–847.

43. Keren-Shaul H, Spinrad A, Weiner A, et al. A unique microglia type associated with restricting development of Alzheimer’s disease. Cell 2017;169(7):1276–1290. e17.

44. Krasemann S, Madore C, Cialic R, et al. The TREM2-APOE pathway drives the transcriptional phenotype of dysfunctional microglia in neurodegenerative diseases. Immunity 2017;47(3):566–581. e9.

45. Fridman AL, Tainsky MA. Critical pathways in cellular senescence and immortalization revealed by gene expression profiling. Oncogene 2008;27(46):5975–87. (In eng). DOI: 10.1038/onc.2008.213.

46. Casella G, Munk R, Kim KM, Piao Y, De S, Abdelmohsen K, Gorospe M. Transcriptome signature of cellular senescence. Nucleic Acids Res 2019;47(14):7294–7305. (In eng). DOI: 10.1093/nar/gkz555.

47. Hernandez-Segura A, de Jong TV, Melov S, Guryev V, Campisi J, Demaria M. Unmasking Transcriptional Heterogeneity in Senescent Cells. Curr Biol 2017;27(17):2652–2660.e4. (In eng). DOI: 10.1016/j.cub.2017.07.033.

48. Bigenwald C, Le Berichel J, Wilk CM, et al. BRAFV600E-induced senescence drives Langerhans cell histiocytosis pathophysiology. Nature Medicine 2021;27(5):851–861.DOI: 10.1038/s41591-021-01304-x.

49. Hu Y, Fryatt GL, Ghorbani M, et al. Replicative senescence dictates the emergence of disease-associated microglia and contributes to Aβ pathology. Cell Reports 2021;35(10):109228. DOI: 10.1016/j.celrep.2021.109228.

50. Wen J, Wang S, Guo R, Liu D. CSF1R inhibitors are emerging immunotherapeutic drugs for cancer treatment. Eur J Med Chem 2023;245(Pt 1):114884. (In eng). DOI:10.1016/j.ejmech.2022.114884.

51. Ribeiro MJ, Idbaih A, Thomas C, et al. 18F-FDG PET in neurodegenerative Langerhans cell histiocytosis : results and potential interest for an early diagnosis of the disease. J Neurol 2008;255(4):575–80. DOI: 10.1007/s00415-008-0751-8.

52. Haroche J, Cohen-Aubart F, Emile JF, et al. Dramatic efficacy of vemurafenib in both multisystemic and refractory Erdheim-Chester disease and Langerhans cell histiocytosis harboring the BRAF V600E mutation. Blood 2013;121(9):1495–500. DOI: 10.1182/blood-2012-07-446286.

53. Martincorena I, Roshan A, Gerstung M, et al. Tumor evolution. High burden and pervasive positive selection of somatic mutations in normal human skin. Science 2015;348(6237):880–6. DOI: 10.1126/science.aaa6806.

54. Martincorena I, Fowler JC, Wabik A, et al. Somatic mutant clones colonize the human esophagus with age. Science 2018;362(6417):911–917. DOI: 10.1126/science.aau3879.

55. Reiner A, Yekutieli D, Benjamini Y. Identifying differentially expressed genes using false discovery rate controlling procedures. Bioinformatics 2003;19(3):368–75. DOI: 10.1093/bioinformatics/btf877.

56. Zehir A, Benayed R, Shah RH, et al. Mutational landscape of metastatic cancer revealed from prospective clinical sequencing of 10,000 patients. Nat Med 2017;23(6):703–713. DOI: 10.1038/nm.4333.

57. Bardou P, Mariette J, Escudie F, Djemiel C, Klopp C. jvenn: an interactive Venn diagram viewer. BMC Bioinformatics 2014;15:293. DOI: 10.1186/1471-2105-15-293.

58. Zhang H, Karnoub ER, Umeda S, et al. Application of high-throughput single-nucleus DNA sequencing in pancreatic cancer. Nat Commun 2023;14(1):749. DOI:10.1038/s41467-023-36344-z.

59. Demaree B, Delley CL, Vasudevan HN, Peretz CAC, Ruff D, Smith CC, Abate AR. Joint profiling of DNA and proteins in single cells to dissect genotype-phenotype associations in leukemia. Nat Commun 2021;12(1):1583. DOI: 10.1038/s41467-021-21810-3.

60. Sashittal P, Zhang H, Iacobuzio-Donahue CA, Raphael BJ. ConDoR: tumor phylogeny inference with a copy-number constrained mutation loss model. Genome Biol 2023;24(1):272. DOI: 10.1186/s13059-023-03106-5.

61. Dobin A, Davis CA, Schlesinger F, et al. STAR: ultrafast universal RNA-seq aligner. Bioinformatics 2013;29(1):15–21. DOI: 10.1093/bioinformatics/bts635.

62. Liao Y, Smyth GK, Shi W. The R package Rsubread is easier, faster, cheaper and better for alignment and quantification of RNA sequencing reads. Nucleic Acids Res 2019;47(8):e47. DOI: 10.1093/nar/gkz114.

63. McCarthy DJ, Chen Y, Smyth GK. Differential expression analysis of multifactor RNA-Seq experiments with respect to biological variation. Nucleic Acids Res 2012;40(10):4288–97. DOI: 10.1093/nar/gks042.

64. Raudvere U, Kolberg L, Kuzmin I, Arak T, Adler P, Peterson H, Vilo J. g:Profiler: a web server for functional enrichment analysis and conversions of gene lists (2019 update). Nucleic Acids Res 2019;47(W1):W191–W198. DOI: 10.1093/nar/gkz369.

65. Parkhurst CN, Yang G, Ninan I, et al. Microglia promote learning-dependent synapse formation through brain-derived neurotrophic factor. Cell 2013;155(7):1596–609. DOI: 10.1016/j.cell.2013.11.030.

66. Srinivas S, Watanabe T, Lin CS, William CM, Tanabe Y, Jessell TM, Costantini F. Cre reporter strains produced by targeted insertion of EYFP and ECFP into the ROSA26 locus. BMC Dev Biol 2001;1:4. DOI: 10.1186/1471-213x-1-4.

67. Stoeckius M, Zheng S, Houck-Loomis B, et al. Cell Hashing with barcoded antibodies enables multiplexing and doublet detection for single cell genomics. Genome Biol 2018;19(1):224. DOI: 10.1186/s13059-018-1603-1.

68. Azizi E, Carr AJ, Plitas G, et al. Single-Cell Map of Diverse Immune Phenotypes in the Breast Tumor Microenvironment. Cell 2018;174(5):1293–1308 e36. DOI: 10.1016/j.cell.2018.05.060.

69. Levine JH, Simonds EF, Bendall SC, et al. Data-Driven Phenotypic Dissection of AML Reveals Progenitor-like Cells that Correlate with Prognosis. Cell 2015;162(1):184–97. DOI: 10.1016/j.cell.2015.05.047.

70. Finak G, McDavid A, Yajima M, et al. MAST: a flexible statistical framework for assessing transcriptional changes and characterizing heterogeneity in single-cell RNA sequencing data. Genome Biol 2015;16:278. DOI: 10.1186/s13059-015-0844-5.

71. Liddelow SA, Guttenplan KA, Clarke LE, et al. Neurotoxic reactive astrocytes are induced by activated microglia. Nature 2017;541(7638):481–487. DOI: 10.1038/nature21029.

72. Habib N, McCabe C, Medina S, et al. Disease-associated astrocytes in Alzheimer’s disease and aging. Nat Neurosci 2020;23(6):701–706. DOI: 10.1038/s41593-020-0624-8.

